# Glial Hedgehog and lipid metabolism regulate neural stem cell proliferation in *Drosophila*

**DOI:** 10.1101/2020.05.18.100990

**Authors:** Qian Dong, Michael Zavortink, Francesca Froldi, Sofya Golenkina, Tammy Lam, Louise Y. Cheng

## Abstract

The final size and function of the adult central nervous system (CNS) is determined by neuronal lineages generated by neural stem cells (NSCs) in the developing brain. In *Drosophila*, NSCs called neuroblasts (NBs) reside within a specialised microenvironment called the glial niche. Here, we explore non-autonomous glial regulation of NB proliferation. We show that lipid droplets (LDs) which reside within the glial niche are closely associated with the signalling molecule Hedgehog (Hh). Under physiological conditions, cortex glial Hh is autonomously required to sustain niche chamber formation, and non-autonomously restrained to prevent ectopic Hh signalling in the NBs. In the context of cortex glial overgrowth, induced by Fibroblast Growth Factor (FGF) activation, Hh and lipid storage regulators Lsd-2 and Fasn1 were upregulated, resulting in activation of Hh signalling in the NBs; which in turn disrupted NB cell cycle progression and reduced neuronal production. We show that the LD regulator Lsd-2 modulates Hh’s ability to signal to NBs, and de novo lipogenesis gene Fasn1 regulates Hh post-translational modification via palmitoylation. Together, our data suggest that the glial niche non-autonomously regulates NB proliferation and neural lineage size via Hh signaling that is modulated by lipid metabolism genes.

## Introduction

Most stem cells reside within specialized groups of cells, collectively referred to as a niche, that provide the trophic, structural and nutritional microenvironment to sustain and protect the stem cells during development (Scadden, 2014). The niche relays developmental and physiological states of the animal to the stem cells and influences the stem cells’ ability to divide in accordance with the environmental state of the organism. Asymmetrically dividing and multipotent neural stem cells in both mammals and invertebrates are responsible for generating the adult nervous system (Homem and Knoblich, 2012).

In *Drosophila*, the vast majority of NBs are specified during embryogenesis, proliferate throughout larval development, and terminate divisions during pupal stages. Type I NBs located within the ventral nerve cord (VNC) and the central brain (CB), are the predominant type of NBs, while type II NBs are eight NB lineages located on the dorsal surface of the central brain (Homem and Knoblich, 2012). During each type I NB cell division, NB self-renews, and produces a smaller ganglion mother cell (GMC) that creates a limited number of neurons or glia. The ability of NBs to divide and generate appropriate progeny number and cell diversity is determined by their ability to maintain asymmetric division, regulate the speed of their cell cycles, and timely enter/exit the cell cycle at the beginning and end of neurogenesis (Homem and Knoblich, 2012). Cell intrinsic mechanisms such as the temporal regulation of NB identity via transcription factors that are expressed throughout the life time of the NBs, impact on both the numbers and the types of neurons generated by the NB (reviewed by Doe, 2017). However, more recently, attention has shifted towards understanding how cell extrinsic signals are interpreted by the NBs to alter their behavior (reviewed by Ramon-Canellas et al., 2019).

Larval NBs and their progeny are surrounded by a scaffold of glial cell processes, which form the stem cell niche in the CNS (Figure 1A). Glial cells fall into three classes: (i) surface (perineural and subperineural) glia that enwrap the CNS to form the blood brain barrier (BBB); (ii) cortex glia that encapsulate neuronal soma and NBs; and (iii) neuropil glia that are located at the cortex–neuropil interface and form a sheath around the neuropil compartments (reviewed by Freeman, 2015).

**Figure 1:**
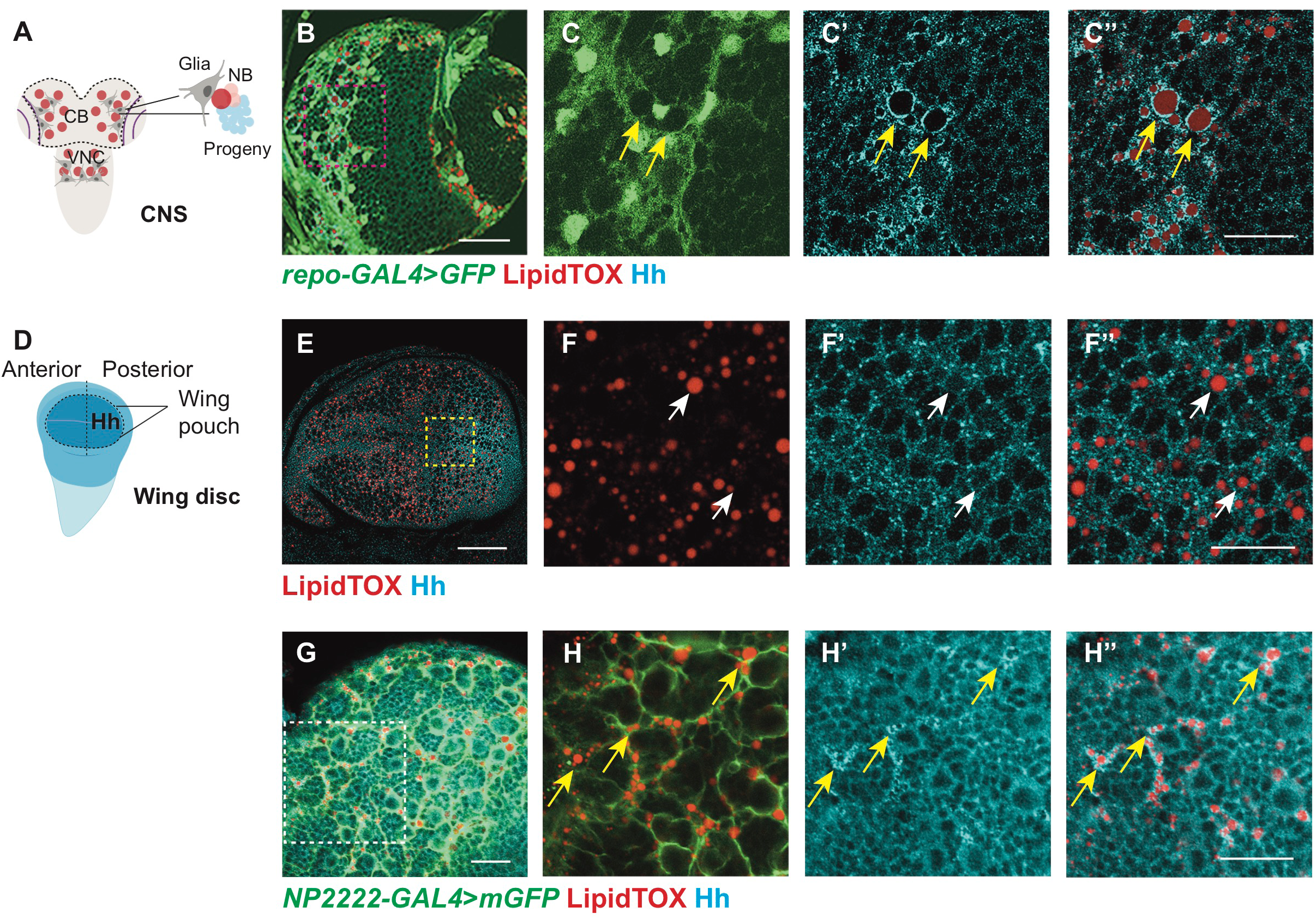
Hh is localized to the LDs within cortex glial cells. Images in this and following figures are of larval central brains at 96ALH, scale bar = 50 μm, in all graphs, error bars showed standard error of the mean (SEM) unless otherwise specified. Data information is described as: Mean ± SEM, n= the number of samples. *(*) p<0.05 (**) p< 0.01, (***) p< 0.001, (****) p < 0.0001, (ns) p>0.05.* A) Schematic depicting developing *Drosophila* CNS, where in the CB, glial niche surrounds NBs and its progeny. B-C’’) Representative images showing Hh accumulates on the surface of LDs in glial cells (*repo-GAL4>GFP*) of the CB (yellow arrows). D) Schematic depicting the developing *Drosophila* wing disc, where Hh is expressed in the posterior compartment. E-F’’) In the posterior compartment of the wing pouch, LDs and Hh are not tightly associated (white arrows). G-H’’) Hh - LD associations are observed in the cortex glia (*NP2222-GAL4>mGFP*) Hh is detected with a Hh antibody and LDs are visualized with LipidTOX unless otherwise stated. C-C’’, F-F’’, H-H’’ are zoomed in images of B, E, G, respectively. Scale bar=20 µm in C-C’’, G-H’’, Scale bar=10 µm in F-F’’.

The intimate relationship between glial cells and NBs has been extensively studied in the context of NB entry into the cell cycle at the beginning of postembryonic neurogenesis shortly after larval hatch (reviewed by Ding et al., 2020). Feeding has been shown to trigger insulin production by surface glial cells, which in turn activates the insulin/insulin-like growth factor pathway in neighboring NBs and stimulates their growth and proliferation via activation of the Phosphoinositide 3-kinase (PI3K) signaling pathway (Chell and Brand, 2010; Sousa-Nunes et al., 2011). Once NBs enter into the cell cycle, glial cells continue to play active roles in promoting NB proliferation. These reactivated NBs are found in close association with cortex glia (Hoyle, 1986; Hoyle et al., 1986; Pereanu et al., 2005), and this contact is maintained through adhesion via E-cadherin. Disruption of NB-cortex glia contact affects the NB’s ability to undergo mitosis (Doyle et al., 2017; Dumstrei et al., 2003), and the failure to expand the glial membrane also affects both neuronal survival as well as NB cell cycle progression (Speder and Brand, 2018; Yuan et al., 2020). Diffusible molecules that pass from glial cells to influence NB behavior include Dally-like (Dlp) in the perineural glia, (Kanai et al., 2018) and Jellybelly (Jeb) in the cortex glia (Cheng et al., 2011). Furthermore, organelles such as lipid droplets (LDs) in the glial niche have been shown to buffer NBs proliferation from peroxidation chain reactions induced by oxidative stress (Bailey et al., 2015), suggesting that glial niche and the signaling molecules produced by these cells are important mediators of non-autonomous regulation of NB behavior during development and environmental stress.

In this study, we investigate how the stem cell niche and its dysfunction influence stem cell behavior and the consequences on the brain as a whole. We found that the signaling molecule Hedgehog (Hh), involved in numerous developmental processes, resides within the cortex glial membrane that surrounds NBs. Hh is autonomously required to promote glial niche growth as well as acts non-cell autonomously to activate the Hh signaling in the NB, triggering its delay in S-phase progression. Maintaining cortex glial size is important, as overgrowth induced by FGF activation, a mutation implicated in glioblastoma (reviewed by Dienstmann et al., 2014; Jimenez-Pascual and Siebzehnrubl, 2019; Morrison et al., 1994; Yamada et al., 1999), phenocopied the effects of glial Hh activation on NBs. Indeed, inhibiting Hh, rescued NB proliferation defects. Furthermore, we demonstrated that downstream of glial FGF signaling, Hh activity and its ability to signal to NBs is modulated by two lipid storage regulators Lsd-2 and Fasn1. Together, our data show that a dysfunctional niche can non-autonomously affect NB’s ability to produce the correct number of neurons that make up the adult CNS. This process mechanistically involves the Hh signaling pathway and its modulation by lipid metabolism.

## Results

### Hh is localised to the LDs within cortex glial cells

To identify potential morphogens that facilitate glia-NB communication in the CB, we assayed for secreted molecules which are known to act in a paracrine fashion. We found that Hh is expressed at high levels at 96 hours After Larval Hatching (96ALH) in glial cells labelled using *Repo-GAL4 > GFP* (Figure 1A-C’). Hh is a morphogen that was first identified to regulate embryo segmentation and wing imaginal disc development (Heemskerk and DiNardo, 1994; Nusslein-Volhard and Wieschaus, 1980). In the wing disc, it is expressed in the posterior compartment and is distributed in a gradient to regulate target gene expression in the anterior compartment (Figure 1D-E; Chen et al., 2017). In the glial niche, however, we found Hh staining was mostly distributed in ring-like structures in the glial cytoplasm (yellow arrows, Figure 1B-C’). We then assessed if Hh is associated with specific organelles. LDs are round-shaped organelles, consisting of a hydrophobic core for the storage of neutral lipids and a phospholipid monolayer containing LD surface proteins. As LDs have previously been reported to be enriched in the glial niche (Bailey et al., 2015; Kis et al., 2015), we therefore tested if Hh ligands are associated with LDs. Using a neutral lipid stain, lipidTOX to visualise LDs and either an antibody, or a BAC encoded Hh:GFP to detect Hh (Chen et al., 2017), we found that Hh is localised to the surface of the LDs in the glial niche (yellow arrows, Figure 1B-C’’Figure S1A-B’’), but not in the wing disc (white arrows, Figure 1E-F’’, Figure S1C-D’’) nor at earlier stages of development in the CB (48 ALH, Figure S2 A-B’’). Together, our data suggests that Hh is localised to the surface of LDs in the glial niche.

**Figure 2:**
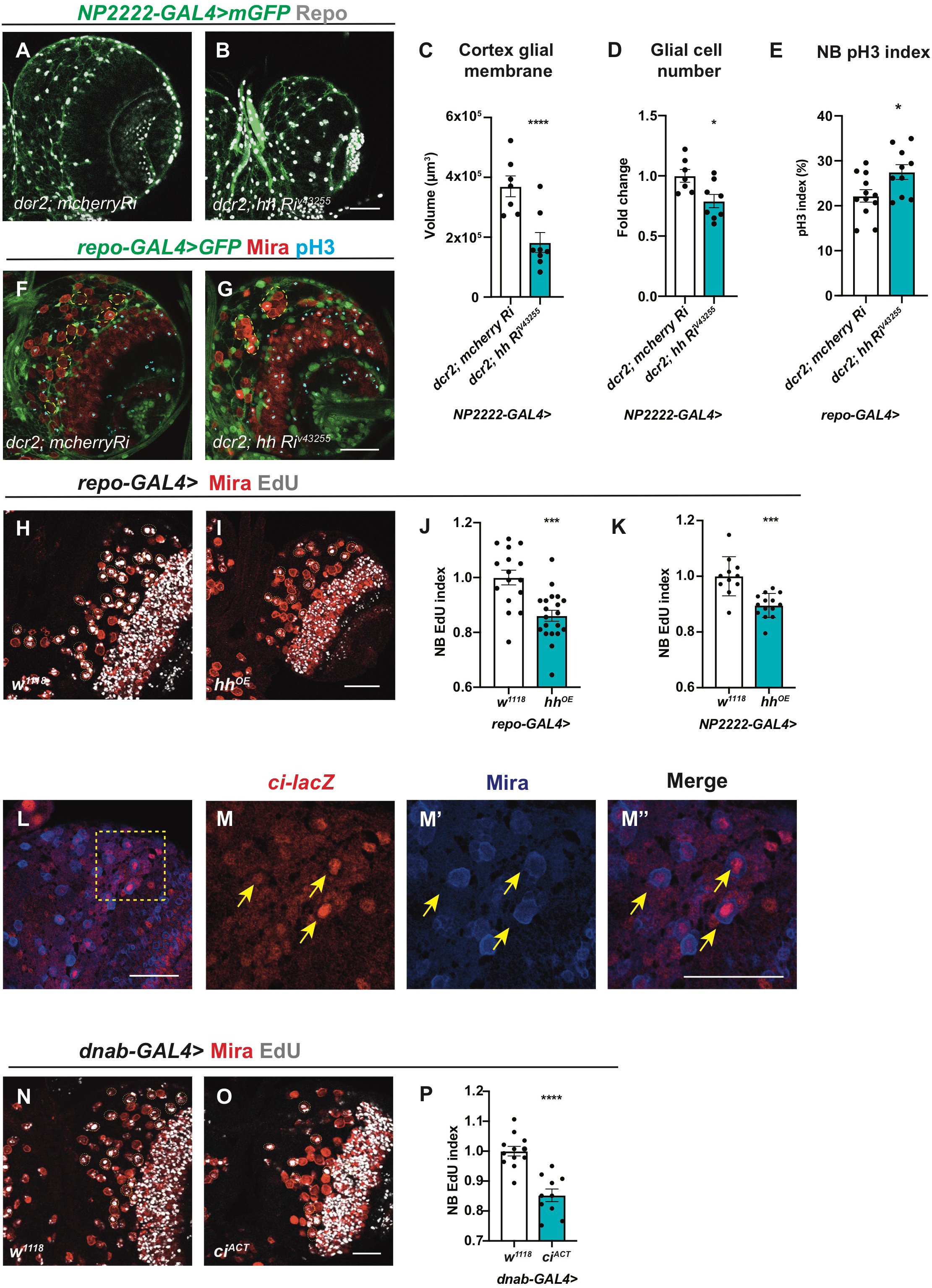
Hh autonomously regulates cortex gliogenesis and non-autonomously regulates NB proliferation. A-D) Representative images showing that upon knockdown of Hh in cortex glial cells (*NP2222-GAL4>mGFP* with *UAS-dcr2*), cortex glial membrane and overall Repo+ glial cell number are significantly reduced, quantified in C and D, respectively. E-G) Hh knockdown in glia (*repo-GAL4>GFP*) results in niche disruption and clustering of NBs (circled with yellow dashed line), as well as an increase in the percentage of NBs in M phase (pH3^+^), quantified in E. H-K) Representative images showing that Hh overexpression using pan-glial (*repo-GAL4*) and cortex-glial (*NP2222-GAL4*) drivers both result in a decrease in NB EdU index, quantified in J and K, respectively. L-M’’) Representative images showing *ci-lacZ* is expressed in NBs (yellow arrows). M-M’’ are zoomed in images of L. N-P) Overexpression of *ci^ACT^* in NBs (*dnab-GAL4*) reduces EdU index, quantified in P. NBs are marked with Mira and EdU^+^ NBs are circled by yellow dashed line.

We next explored whether Hh-LD associations are specifically localised to a glial subtype. Hh-LD associations were largely absent from both surface glial cells that forms the blood brain barrier (BBB) of the CNS (white arrows, Figure S2E-F’’’), as well as optic lobe glial cells (white arrows, Figure S2C-D’’’). In fact, Hh-LD associations were enriched in the cortex glial cells, underneath the sheath-like surface glial clone generated via *repo*-MARCM (yellow arrows, Figure S2G). Using a cortex specific driver, *NP2222-GAL4* (Awasaki et al., 2008; Hayashi et al., 2002), we confirmed that the Hh-LD associations were localised to the cortex glia (Figure 1G-H’’).

### Hh autonomously regulates cortex gliogenesis and non-autonomously regulates NB proliferation

In the mouse brain, Hh has been shown to promote astrocyte proliferation (Chandra et al., 2015; Takezaki et al., 2011; Ugbode et al., 2017). In glioblastoma (GBM), the Hh/Gli1 signaling pathway acts to accelerate cell proliferation (Chandra et al., 2015). To investigate the role of Hh in *Drosophila* cortex glial cells, where Hh is most abundant, we used cortex specific *NP2222-GAL4* to express *hh* RNAi together with *UAS-Dcr2* (*Dicer-2*) which sufficiently depleted Hh expression (Figure S3A-B) and reduced overall CNS size. The reduction in CNS volume was accounted for by a significant decrease in Repo^+^ glial cell number and membrane size (labelled using *NP2222-GAL4>GFP*, Figure 2A-D). Using a pan-glial driver *repo-GAL4*, we found cortex glial chambers were significantly disrupted upon Hh knockdown (Speder and Brand, 2018; Yuan et al., 2020), leading to clustering of NBs (compare Figure 2G to F). However, overexpression of Hh did not cause glial overgrowth (data not shown). This suggests Hh is necessary but not sufficient for glial expansion during CNS development.

Given the role of Hh as a secreted ligand that can act over short range within the NB lineage (Chai et al., 2013) and that it is highly expressed in the cortex glial niche surrounding NBs, it is plausible that glial Hh non-autonomously affect NB proliferation. We next explored the potential impact of glial Hh on the activities of Type I NBs. As Hh is required to maintain the glial niche (Figure 2 A-D, F-G) and niche impairment has been shown to induce NB elimination (Read, 2018), we first assessed for alterations in NB number. We found that pan-glial Hh knockdown (*repo-GAL4*) did not significantly alter NB number (Figure S3E), suggesting that NB survival is unaffected. We then investigated the effects of glial Hh knockdown on NB proliferation. Glial Hh knockdown using *repo-GAL4*, induced a small increase in the percentage of NBs in M phase (pH3 index; Figure 2E). To assess S phase progression, we assessed EdU (5-ethynyl-2′-deoxyuridine) incorporation during a 15-minute time window (EdU index, Figure S3D, yellow arrows). Here, we found EdU incorporation was significantly reduced upon glial Hh knockdown, suggesting that fewer NBs entered into the S phase of the cell cycle (Figure S3C). Interestingly, a similar NB phenotype was reported by Speder and Brand, 2018, caused by niche impairment induced via overexpression of glial ΔP60 (which inhibits PI3K signalling pathway). Therefore, our data suggests that the cell cycle defects we observed could be a result of glial niche impairment.

The sub-perineural glial Dlp and the cortex glial Jeb promote NB proliferation during development; however, overexpression of these signaling molecules in the glial niche was not sufficient to increase NB cell cycle rate (Cheng et al., 2011; Kanai et al., 2018). We next assessed the effect of glial Hh overexpression on NB behaviour. Pan-glial induction of Hh, did not significantly alter NB number (Figure S3E). However, pan-glial and cortex glial specific Hh overexpression caused a reduction in NB EdU incorporation (Figure 2H-K). Using reporter lines of Hh activity *ci* -*lacZ* (Schwartz et al., 1995) and Ptc:mCherry (Chen et al., 2017; Varjosalo and Taipale, 2008), we found that Hh signaling is highly active in the NBs (Figure 2L-M’’, Figure S3F-G’’), consistent with a previous report by Chai et al., 2013. Furthermore, activation of Hh transcriptional activator *cubitus interruptus* (*ci^Nc5m5m^* or *ci^ACT^*) with a NB specific driver *dnab-GAL4* (Maurange et al., 2008) significantly reduced NB EdU index (Figure 2N-P), phenocopying the effects of glial-Hh overexpression. Together, our data suggests that high levels of glial Hh expression restricts NB cell cycle progression.

### Hh activity is modulated by Lsd-2

Lipid storage droplet-2 (Lsd-2) is the *Drosophila* orthologue of the mammalian perilipin2, and a widely used marker of LDs (Teixeira et al., 2003, Figure 3 A-B’’). Using a GFP reporter against Lsd-2 together with Hh antibody, we observed that Lsd-2 and Hh colocalize to the surface of LDs in the cortex glia (yellow arrows, Figure 3C-D’’). To explore whether Lsd-2 affects Hh activity, we knocked down Lsd-2 while overexpressing Hh in cortex glial cells (*NP2222-GAL4>*). Lsd-2 is known to block the access of lipases, thus promoting lipid storage (Teixeira et al., 2003). As expected, knockdown of Lsd-2 caused a significant reduction in LDs (Figure 3E-G). Furthermore, this also significantly rescued the NB EdU incorporation defects caused by *hh* overexpression (Figure 3H-K), suggesting Lsd-2 modulates Hh’s ability to signal to NBs.

**Figure 3:**
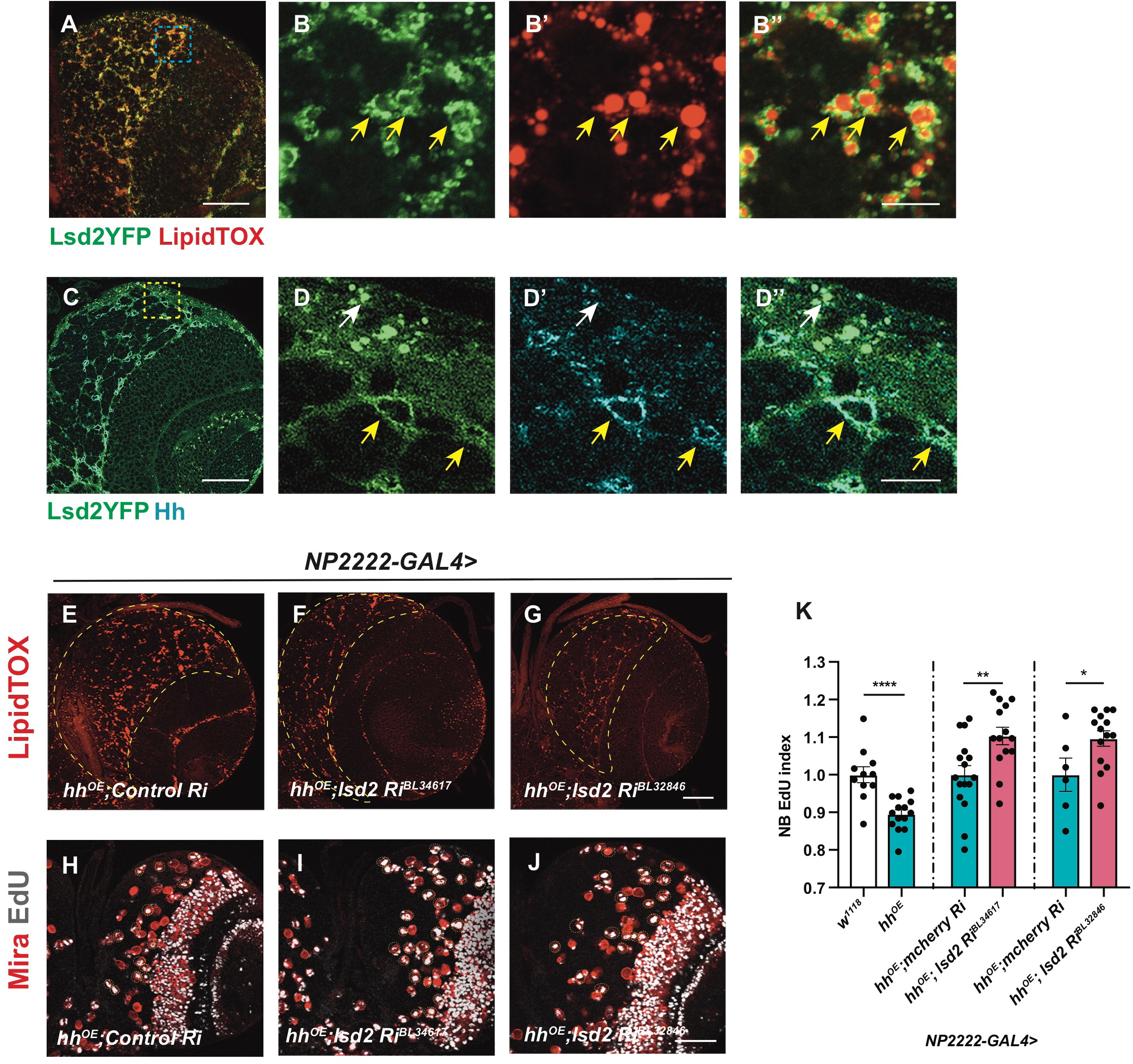
Hh activity is modulated by Lsd-2. A-B’’) Representative images showing that Lsd2YFP is localised to the surface of LDs (yellow arrows). C-D’’) Lsd2YFP co-localises with Hh antibody staining in the cortex glia (yellow arrows) but not surface glial cells (white arrow). E-G) Lsd-2 knockdown in cortex glial cells (*NP2222-GAL4*) where *hh* is overexpressed effectively reduces LD number in CB (outlined in yellow dashed lines). H-K) Representative images showing that NB EdU index is rescued upon Lsd-2 knockdown in cortex glial cells (*NP2222-GAL4*) where *hh* is overexpressed, quantified in K. K), first column depicts the same data as Figure 2K. B-B’’, D-D’’are zoomed in images of A and C. Scale bar=10 µm in B-B’’ and D-D’’.

### FGFR, but not EGFR or InR, activation induces cortex glial overgrowth

Given that we found Hh activity is modulated by a lipid modulator Lsd-2, it is possible that Hh might act in disease contexts such as glioblastoma, where glial cells undergo increased proliferation and lipid metabolism alterations (Geng and Guo, 2017). As Hh/LD associations mostly localised to the cortex glia that enwraps NBs, we explored the role for cortex glial overgrowth, using previously characterized glioma models, involving the activation of FGF, InR and EGFR (Avet-Rochex et al., 2012; Read et al., 2009; Reddy and Irvine, 2011; Witte et al., 2009). Firstly, we characterised the effect of overexpression of a wild type form of InR, constitutively activated form of the FGF receptor called Heartless (Htl) and EGFR. We found *htl^ACT^* but not *InR^wt^* or *Egfr^ACT^* overexpression caused an expansion of the cortex glial niche which enwraps NBs (Figure 4A-D’). This observation was further confirmed by driving these transgenes with cortex glial specific *NP2222-GAL4*. As overexpression of *Egfr^ACT^* using *NP2222-GAL4* caused lethality, an alternative cortex glial driver, *wrapper-GAL4* (Coutinho-Budd et al., 2017; Richier et al., 2017) was utilized. Analysis of the total glial cell number indicates overexpression of *htl^ACT^*, but not *InR^wt^* or *Egfr^ACT^*, led to an increase in the number of cortex glial cells (Figure 4E-G, I-K). Furthermore, *htl^ACT^* overexpression significantly increased the size of the cortex glial membrane (Figure 4E-H). Together, our data suggest the activation of FGF, but not InR or EGFR, induced cortex glial niche overgrowth.

**Figure 4:**
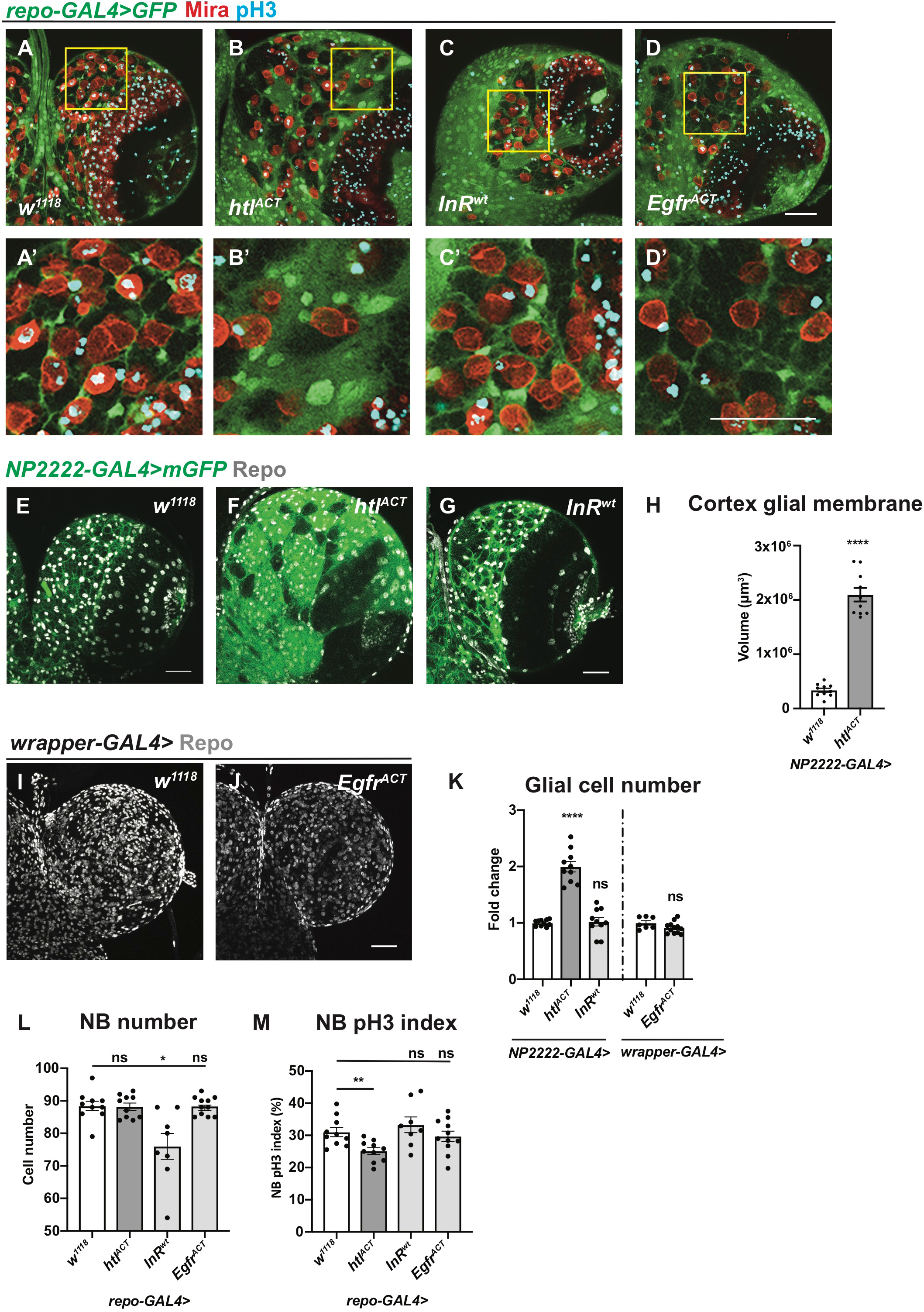
Activation of *htl^ACT^* but not *InR^wt^* and *Egfr^ACT^* induces cortex glial overgrowth. A-D’) Representative images showing that pan-glial overexpression of *htl^ACT^*, but not *InR^wt^* or *Egfr^ACT^* causes an expansion of cortex glia that enwraps NBs. Glial cells are marked with *repo-GAL4 >GFP*, and NBs are marked with Mira. A’, B’, C’ and D’ are zoomed in images of A, B, C and D, respectively. E-K) Representative images showing that cortex glial overexpression of *htl^ACT^* but not *InR^wt^* or *Egfr^ACT^* causes an increase in glial cell (Repo^+^) numbers and cortex glial membrane size, quantified in K and H. *NP2222-GAL4>mGFP* is used to mark cortex glial membrane in E-G and *wrapper-GAL4>* is used in I-J. L) Glial (*repo-GAL4>*) overexpression of *InR^wt^* but not *htl^ACT^* or *Egfr^ACT^* significantly reduces the number of CB NBs. M) Glial (*repo-GAL4>*) overexpression of *htl^ACT^* but not *InR^wt^* or *Egfr^ACT^* significantly reduces the pH3 index of CB NBs (marked with Mira and pH3 in A-D’).

We next explored the impact of glial niche overgrowth on NBs. Using the pan-glial driver *repo-GAL4*, we measured NB cell number and its impact on the cell cycle assessed with pH3 index. We found that *InR^wt^*, but not *htl^ACT^* and *Egfr^ACT^* overexpression, caused a reduction in NB number, reminiscent of the NB elimination phenotype reported for glial *Pvr^ACT^* overexpression (Read, 2018, Figure 4L). In addition, only overexpression of *htl^ACT^*, which induced cortex glial overgrowth, caused a reduction of NB pH3 index; while *InR^wt^* and *Egfr^ACT^* overexpression did not significantly alter NB cell cycle progression (Figure 4M). Together, these data indicate that cortex glial niche is the key glial subset that mediates glia-NB crosstalk, and its expansion driven by FGF activation affected NB cell cycle progression.

### Cortex glial overgrowth mediated by FGF activation slows down NB cell cycle progression

To further investigate the effect of glial FGF activation on NB cell cycle, we assessed S-phase progression via EdU incorporation. FGF activation using pan glial (*repo-GAL4*) as well as cortex glial (*NP2222-GAL4*) drivers caused a significant reduction in NB EdU index (Figure 5A-E). Consistent with the reduction in pH3 index (Figure 4M), these experiments confirm that the cortex glia is responsible for NB cell cycle delay induced by FGF activation.

**Figure 5.**
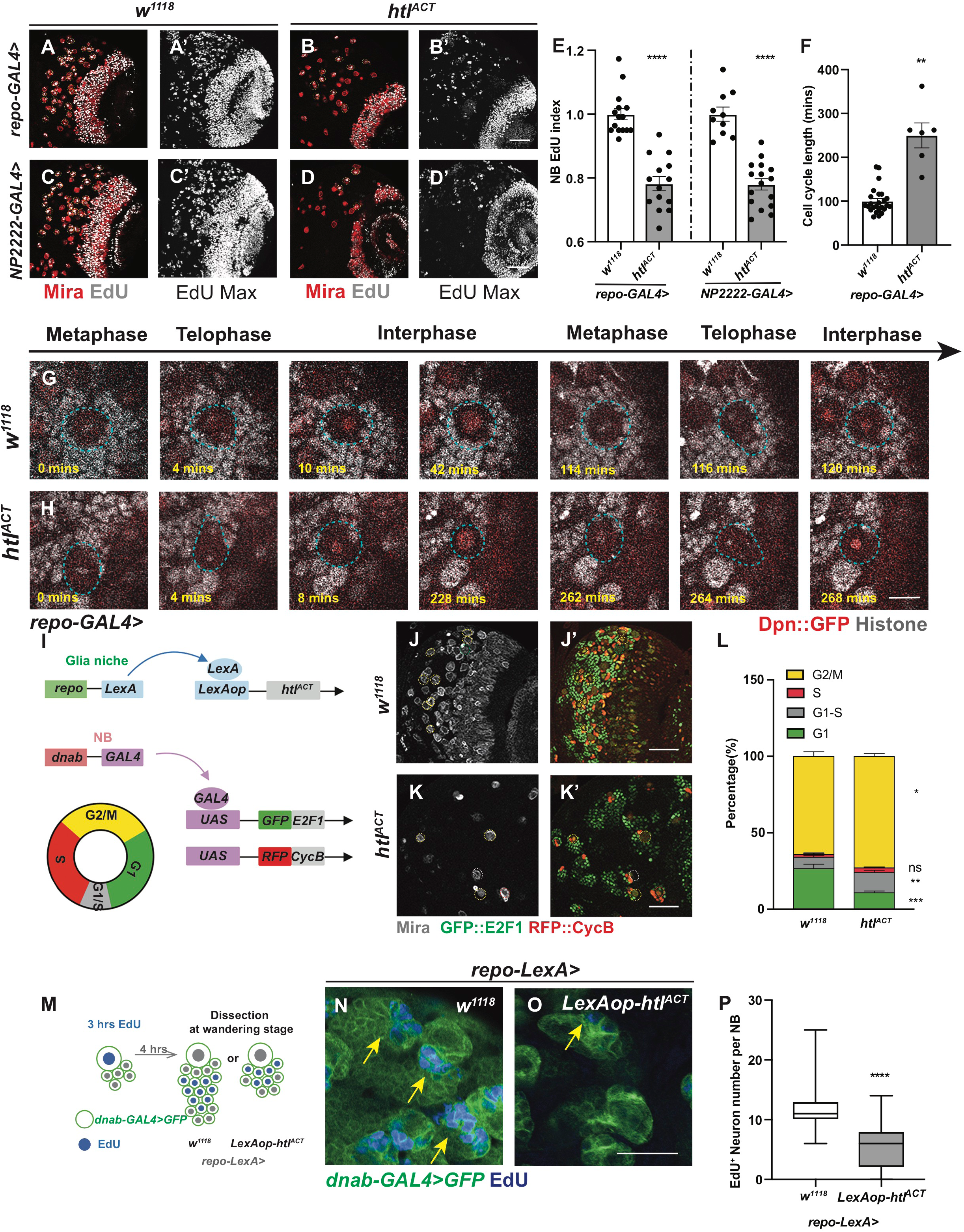
Cortex glial overgrowth mediated by *htl^ACT^* overexpression triggers NB cell cycle delay. A-E) Representative images showing that both pan-glial (*repo-GAL4>*) and cortex glial (*NP2222-GAL4>) htl^ACT^* overexpression significantly reduce NB EdU index, quantified in E. A, B, C, D are single sections with Mira and EdU staining, and A’, B’ C’, D’ are Z-projection of the EdU staining. F-H) Representative still images from ex vivo CNS live imaging at 72ALH showing that pan-glial (*repo-GAL4*) *htl^ACT^* overexpression lengthens NB cell cycle, quantified in F. The cell cycle length is measured as the length between consecutive divisions. NBs (Dpn::GFP, red; Histone RFP, grey) are circled with blue dashed lines. I) Schematic depicting concurrent glial FGF activation (*repo-LexA* >*LexAop-htl^ACT^*), and NB overexpression of fly-FUCCI (*dnab-GAL4>UAS-GFP::E2F1, UAS-RFP::CycB*). The fly-FUCCI system utilizes the fusion protein GFP::E2F1 (a marker for cells in G2, M and G1phase) and RFP::CycB (a marker for cells in S, G2, M phase) to monitor cell cycle progression. Cells in G1 phase are GFP^+^ RFP^-^ (green), cells in G2/ M phase are GFP^+^RFP^+^ (yellow), and cells in S phase are GFP^-^RFP^+^, whereas, cells in G1-S transition are weakly labelled by both GFP and RFP (grey). J-L) Representative images showing that the percentage of NBs in G1-S transition and G2/M phase are both significantly increased with significantly less cells remaining in G1 phase, quantified in L. NBs in G1 phase (Mira^+^, GFP^+^) are circled by green dashed lines; NBs in G1-S transition (Mira^+^, GFP^-^RFP^-^) are circled by grey dashed lines; NBs in S phase (Mira^+^, RFP^+^) are circled by red dashed lines; and NBs in G2/M phase (Mira^+^, GFP^+^RFP^+^) are circled by yellow dashed lines. Statistics: Repo-gal4 > *w^1118^*: G1 (26.72% ± 2.79%, n = 9), G1-S transition (7.36% ± 1.18%, n = 9), S (1.99% ± 0.69%, n = 9), G2/M (63.93% ± 1.76%, n = 9); Repo-gal4 > *htl^ACT^* : G1(10.93% ± 1.02%, n = 10), G1-S transition (13.20% ± 1.43%, n = 10), S(3.08 % ± 0.38%, n = 10), G2/M(72.80% ± 1.76%, n = 10). M) Schematic depicting EdU pulse-chase experiment. Larvae are fed with EdU-containing food for 3 hours and then chased with EdU-free food for 4 hours before CNS dissection at wandering stage. N-P) Representative images showing that the number of EdU^+^ neurons generated per NB is significantly reduced upon pan-glial overexpression of FGF (*repo-LexA > LexAop-htl^ACT^*; yellow arrows), quantified in P (Box plot, Min-Max). NB lineages are marked with *dnab-gal4 > GFP*. Statistics: *repo-LexA> w^1118^*: 11.73 ± 0.32, n = 94 (NB lineages from 5 brain lobes are plotted); *repo-LexA> htl^ACT^*: 5.40 ± 0.33, n =127 (NB lineages from 7 brain lobes are plotted). Scale bar=10 µm in G-H; Scale bar=20 µm in N-O.

We next examined the cell cycle length of NBs using live-cell imaging on explanted brains, where three brains containing multiple NBs labelled with Dpn::GFP and His::RFP (Figure 5G-H) were imaged. We found FGF activation with *repo-GAL4* induced a severe slowing down of the cell cycle at 96ALH, such that we could not capture any entire NB cell cycles within an 8-hour time window. Previously it was reported that NBs cycle faster during earlier developmental stages (Chai et al., 2013; Maurange et al., 2008), and therefore we imaged NB divisions at 72 ALH. We observed that glial FGF activation lengthened NB cell cycle from 100.2±6.0 mins to 250.0±28.4 mins (Figure 5F, Movies S1 and S2).

To decipher which phase of the cell cycle was affected, we generated flies expressing *htl^ACT^* downstream of a *LexA* operator (*LexAop*, Lai and Lee, 2006). Overexpressing *htl^ACT^* using *repo-LexA/LexAop* system enabled us to concurrently induce NB-specific expression of Fly-Fucci using *dnab-GAL4* (as depicted in Figure 5I schematic). With Fly-Fucci, cell cycle phases can be identified using combinations of two fluorescent fusion proteins (Zielke et al., 2014). Surprisingly, we found glial FGF activation significantly increased the percentage of NBs in G2/M and G1-S transition at the expense of cells in G1 phase (Figure 5J-L). As the percentage of NBs in M phase (reflected by pH3 index) was reduced upon glial FGF activation, we conclude that these NBs are potentially stalled at G2/ G2-M and G1-S transitions of the cell cycle. As a result, they cannot efficiently enter S phase or M phase, as indicated by reduced EdU and pH3 indices (Figure 5A-E, Figure 4M), and therefore, undergo a dramatically lengthened cell cycle (Figure 5F).

We expect that slowing down of the NB cell cycle upon glial FGF activation would consequentially affect the number of neurons generated by NBs. To test this hypothesis, we conducted an EdU pulse-chase assay, where the larvae were fed EdU^+^ food for three hours and chased for four hours in EdU-free food (Figure 5M). We then used *dnab-GAL4::UAS-GFP* to mark NB lineages and counted the number of EdU^+^ neurons per lineage. We found that overexpression of *htl^ACT^* with *repo-LexA/LexAop* significantly reduced the number of EdU^+^ neurons generated per NB (Figure 5N-P). Together, our data indicate that FGF activation in cortex glial cells prevents NB cell cycle progression and its ability to produce the correct number of neurons.

### Glial FGF activation affects NB asymmetric division, size and cell cycle exit

Given that cell polarity that contributes to NB asymmetric division is established in the G2/M phase, we then assessed whether NB asymmetric division is affected upon glial FGF activation. In the wildtype, Inscuteable, an adaptor protein that connects the aPKC/Par3/Par6 complex to the PINS/MUD/Gαi complex, forms a crescent at the apical side during NB mitosis (Doe, 1996); these apical complexes further direct the localization of cell fate determinants (Brat/Pros/Numb) and their adaptor proteins Mira (Shen et al., 1997) and PON to the basal cortex (as depicted in Figure S4A schematic). The correct distribution of polarity proteins ensures the generation of a larger daughter NB and a smaller GMC upon asymmetric division (Figure S4H). We found mitotic NBs displayed cytoplasmic or cortical localization of Mira and Insc upon glial FGF activation (Figure S4B-G). Furthermore, telophase NBs and GMCs were also found to be more similar in size (evaluated as NB /GMC size ratio in Figure S4I-K). Intriguingly, both NB and its daughter cells were larger than their wildtype counterparts, consistent with this, the average size of mitotic NBs was also significantly larger upon glial *htl^ACT^* overexpression (Figure S4L, from 12.44±0.16µm to 15.55±0.34µm). This increase in cell size was coupled with elevated cellular growth, as indicated by an increase in nucleoli size (Figure S4M-O, from 1.28±0.05µm to 1.61±0.05µm). It is therefore likely that the delay between consecutive cell cycles allowed these NBs to grow larger.

To investigate when glial FGF starts to impact on NB proliferation, we assessed NB EdU index at 26ALH, a time point when most NBs reactivate from quiescence, and commence post-embryonic neurogenesis (Figure S4P). We found glial *htl^ACT^* overexpression did not significantly affect NB EdU incorporation during a 1-hour EdU pulse (Figure S4Q-S), suggesting that glial FGF does not affect NB reactivation at the beginning of neurogenesis. We then examined whether NB termination at 24 hours after pupal formation (24APF) is altered (Figure S4T). Glial *htl^ACT^* overexpression resulted in the presence of increased number of NBs at 24APF (Figure S4U-W), suggesting NB cell cycle exit is possibly delayed. However, it is not clear whether NBs that persist are capable of dividing. Together we conclude glial-FGF mostly exerts its effects on reducing NB cell proliferation during late larval neurogenesis, coinciding with the time when Hh-LD associations become highly enriched in the cortex glia. Taken together, our results revealed that NB activities including its proliferation, asymmetric division and termination are affected by cortex glial niche overgrowth driven by FGF activation.

### Hh mediates the effects of glial FGF signalling on NB proliferation

Given that glial Hh overexpression or activation of the Hh signalling cascade in NBs similarly inhibits NB proliferation and induces polarity protein delocalisation (Chai et al., 2013); it is therefore plausible that Hh lies downstream of glial FGF to regulate NB activity. To test this hypothesis, we first assessed whether niche derived signals including Hh and its modulators such as LD synthesis enzymes were altered. By RT-qPCR, we found *hh* mRNA was upregulated by 8-fold (normalized to *rpl32*) upon glial FGF activation (*repo-GAL4*, Figure 6F). Amongst genes that promote LD storage (Figure 6I), *fatty acid synthase-1* (*fasn1,* Smith et al., 2003), and *lsd2* (an antagonist of lipases which we have shown to regulate Hh activity, Figure 3), were also significantly upregulated (Figure 6F). Furthermore, Hh staining was found to be more widely distributed in the glial cytoplasm, rather than restricted to the ring-like structures surrounding LDs upon FGF activation (Figure 6A-E, evaluated by the ratio of glial cytoplasm that contains Hh staining).

**Figure 6.**
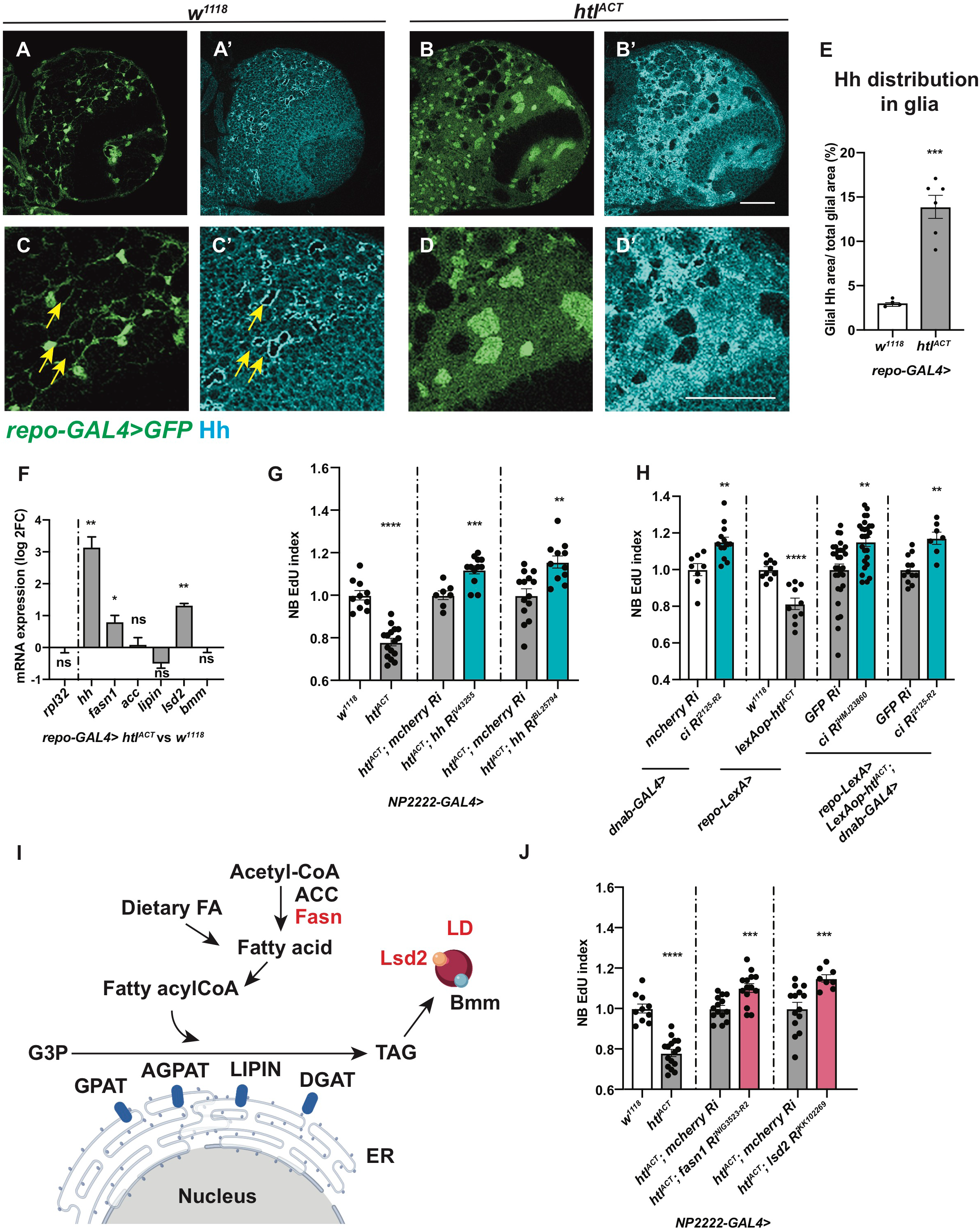
Hh and lipid metabolism regulators mediate the effects of glial *htl^ACT^* on NB proliferation. A-E) Representative images showing that Hh staining normally localised to a ring like structure (yellow arrows), becomes delocalises to the glial cytoplasm upon *htl^ACT^* overexpression, quantified in E. Glial cells are marked with *repo-GAL4>GFP*. C-D’ are zoomed in images of A-B’. F) Pan-glial (*repo-GAL4*) *htl^ACT^* overexpression causes upregulation of *hh, fasn1* and *lsd2* transcripts. The reference gene *rpl32*, lipogenesis genes (*acc*, *lipin*), and lipolysis gene *bmm* transcripts are not significantly altered. The data are represented by log2 fold change relative to the control (*repo-GAL4> w^1118^*). Statistics: *hh:* 3.17 ± 0.33, n = 3; *fasn1*: 0.80 ± 0.20, n = 3; *lsd2:* 1.32 ± 0.06, n = 3. G) Cortex glial (*NP2222-GAL4>*) overexpression of two independent *hh* RNAis significantly rescue EdU incorporation defects caused by *htl^ACT^* overexpression. The *NP2222-GAL4> w^1118^* vs *htl^ACT^* columns depict the same data as those in Figure 5 E. H) Knockdown of NB Hh signalling pathway (*dnab-GAL4> UAS-ciRNAi*) rescues NB EdU incorporation defects induced by glial *htl^ACT^* overexpression (*repo-LexA> LexAop*-*htl^ACT^*). Induction of *ciRNAi* in NBs alone increases NB EdU incorporation. I) Schematic depicting lipogenesis and lipolysis. Lipogenesis begins with *de novo* synthesis of fatty acids by carboxylation of cytosolic acetyl-CoA via acetyl-CoA carboxylase (ACC) and elongation of fatty chain via fatty acid synthase (Fasn, red). Dietary-derived and *de novo-*generated fatty acids are converted into fatty acylCoA, which re-localizes to ER and participates in triglyceride (TAG) synthesis with glycerol-3 phosphate (G3P). This process is mediated by a series of enzymes: glycerol-3-phosphate acyltransferase (GPAT), acylCoA acylglycerol-3-phasphte acyltransferases (AGPAT), Lipin (a phosphatidate phosphatase), diacylglycerol acyltransferase (DGAT, red). TAG is translocated from the ER to the core of the intracellular organelles called LDs. On the surface of LDs, a triglyceride lipase called Brummer (Bmm), and its inhibitor Lsd-2, antagonistically control TAG storage. J) The NB EdU incorporation defects due to cortex glial (*NP2222-GAL4>*) overexpression of *htl^ACT^*, is significantly rescued by overexpression of RNAis against *fasn1* and *lsd2*, compared to corresponding control RNAis. The *NP2222-GAL4> w^1118^* vs *htl^ACT^* columns depict the same data as those in Figure 5 E. The control column for *htl^ACT^; lsd2 Ri^KK102269^* depicts the same data as the control column for *htl^ACT^; hh* Ri*^BL25794^* in Figure 6G.

We next assessed the role of Hh signalling downstream of glial FGF activation. Hh knockdown (RNAi efficiency tested in Tian et al., 2015) rescued NB S-phase delay (Figure 6G) without affecting glial niche size (Figure S5A-B). In addition, we used a *LexA/LexAop* binary expression system in conjunction with the *GAL4/UAS* system, to simultaneously activate glial FGF and inhibit NB Hh signaling. Induction of RNAi against *ci* in the NB caused a significant increase in EdU incorporation (Figure 6H, *dnab-GAL4> ci RNAi*). Together with glial-FGF activation, Ci knockdown also significantly rescued NB EdU index (Figure 6H). Together, our results suggest that Hh mediates glia-NB crosstalk downstream of glial FGF activation.

### Lipid metabolism genes lie downstream of glial FGF-NB crosstalk

As we previously showed that Lsd-2 modulates Hh activity (Figure 3), we hypothesise that lipid metabolism enzymes function upstream of Hh to regulate NB behaviour. Consist with this, we found induction of *hh RNAi* upon cortex glial FGF activation did not alter the number of LDs (Figure S5C-D). We next explored whether lipid metabolism genes mediate the effects of glial FGF activation on NB proliferation. Glial expression of RNAis targeting lipogenesis genes *fasn1* and *lsd2* efficiently reduced LDs (Figure S5E-G), but were not required for wildtype NB S phase progression (Figure S5 H). However, knockdown of Fasn1 and Lsd2 significantly rescued NB S-phase progression defects caused by *htl^ACT^* overexpression (Figure 6J, Figure S5I), suggesting lipid metabolism genes function downstream of glial FGF-NB crosstalk.

### Fasn1 and Rasp affect glial Hh palmitoylation to regulate NB cell cycle

Hh is known to be synthesized as a precursor protein (HhN) which undergoes a series of post-translational modifications within the secretory pathway (Figure 7A; reviewed by Mann and Beachy, 2004). The N- and C-termini of Hh proteins are covalently modified with palmitate and cholesterol, respectively (Pepinsky et al., 1998; Porter et al., 1996). Palmitate is a 16-carbon saturated fatty acid, which can either be diet-derived or synthesized via de novo fatty acid synthesis, mediated by enzymes such as Fasn and ACC (Innis, 2016; Schiller and Bensch, 1971; Slakey et al., 1979). Given Fasn1, an enzyme involved in de novo synthesis of palmitate, is also involved in glial-FGF-NB crosstalk, we assessed whether Hh palmitoylation is required downstream of glial FGF signaling. Palmitoylation of the N-terminal of HhN is mediated by a dedicated acyltransferase in the ER called Rasp in *Drosophila* (Figure 7A). In the embryo, Rasp has been shown to be required for Hh diffusion (Chamoun et al., 2001; Lee and Treisman, 2001; Micchelli et al., 2002). We found glial knockdown of Rasp while not required for NB S phase progression (Figure 7B), was sufficient to rescue NB EdU incorporation defects caused by FGF activation (Figure 7B, *NP2222-GAL4*> *htl^ACT^*).

**Figure 7.**
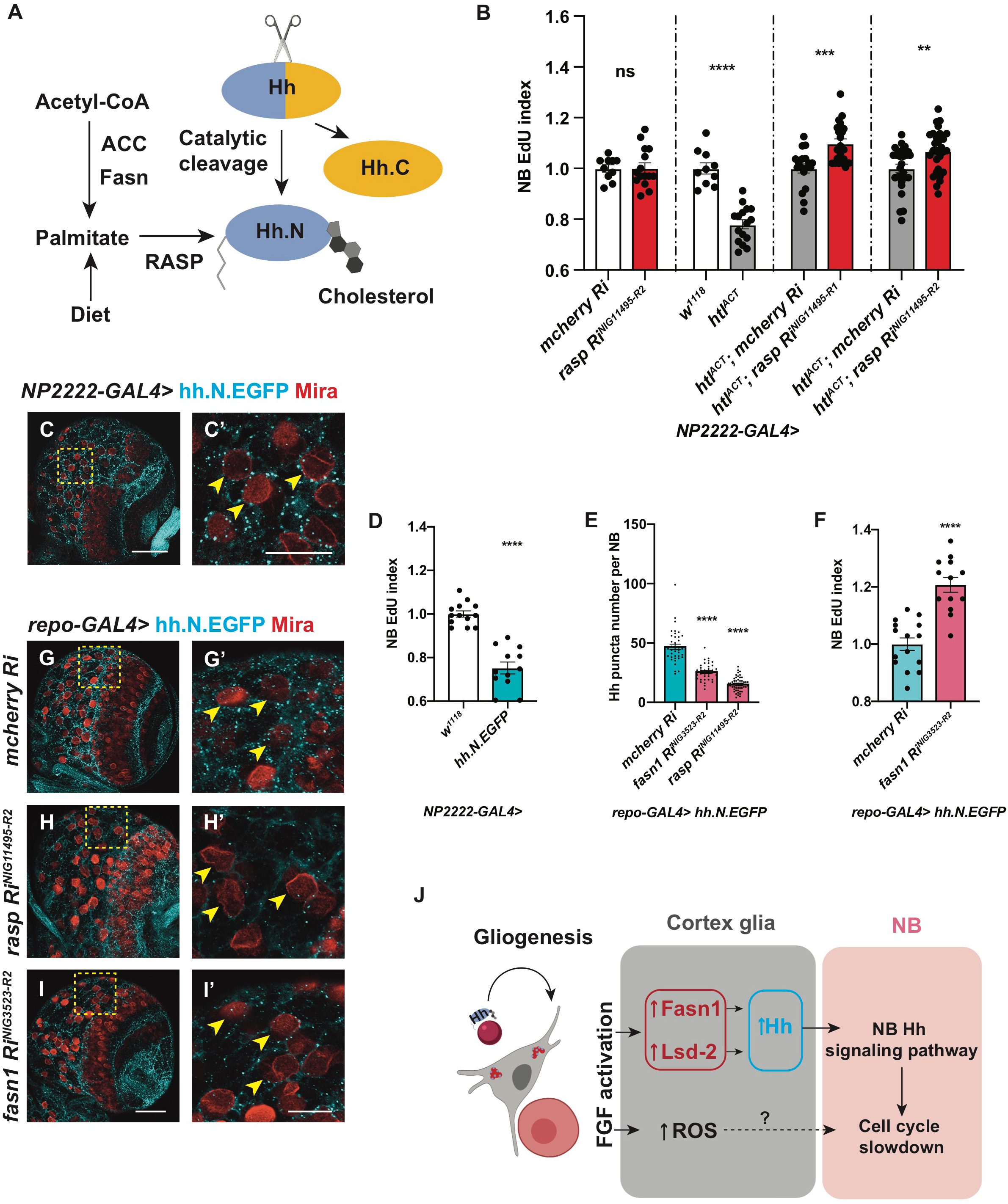
Fasn1 affects glial Hh palmitoylation to regulate its signalling to NBs. A) Schematic depicting Hh auto-processing, which starts with the cleavage of the protein into a C-terminal part (Hh.C, yellow) and a N-terminal part (Hh.N, blue), with simultaneous covalent addition of cholesterol. Palmitate, from either diet or *de novo* lipogenesis (via ACC and Fasn), is added onto Hh-N, in a reaction catalysed by an acyltransferase, encoded by *rasp.* B) Inhibition of palmitoylation (via two independent *rasp RNAis*) rescues NB EdU incorporation defects induced by cortex glial (*NP2222-GAL4>*) *htl^ACT^* overexpression, while knockdown of Rasp in cortex glial cells alone does not alter NB EdU index. The *NP2222-GAL4> w^1118^* vs *htl^ACT^* columns depict the same data as those in Figure 5 E. C-C’) Representative images showing that Hh.N.EGFP (which cannot undergo cholesterol modification) are found as puncta on the surface of NBs (yellow arrows) when overexpressed in neighbouring cortex glial cells (*NP2222-GAL4>*). D) Cortex glial (*NP2222-GAL4>*) overexpression of Hh.N.EGFP significantly reduces NB EdU incorporation. E-I’) Representative images showing that knockdown of Fasn1 in glial cells (*repo-GAL4>*), where Hh.N.EGFP is overexpressed, significantly reduces the number of Hh.N.EGFP puncta on the surface of NBs (yellow arrows), phenocopying the effect of Rasp knockdown, quantified in E. This manipulation also rescues NB EdU incorporation defects, quantified in F. C’, G’, H’ and I’ are zoomed in images of C, G, H and I, respectively; NBs are marked with Mira. J) Schematic depicting our working model. During development, Hh tethered to LDs are localised to cortex glial cells, to activate gliogenesis. Non-autonomously, excessive glial Hh inhibits NB cell cycle progression. Upon cortex glial specific FGF activation, increased Hh modified by Fasn1 and Lsd-2 together with increased ROS prevent NB proliferation. Scale bar=20 μm in C’, G’, H’ and I’.

To test if Hh palmitoylation is required for its ability to signal to NBs, we expressed a GFP tagged and non-cholesterol modifiable form of HhN in cortex glial cells (Hartman et al., 2013; Wendler et al., 2006). HhN-GFP were mostly localised to puncta in the glial cytoplasm, and some of these puncta made contact with NBs (Figure 7C-C’, *NP2222-GAL4*). The number of puncta making contact with NBs was significantly reduced upon knockdown of Rasp (Figure 7H-H’ compared to 7G-G’), suggesting that palmitoylation is required for HhN transport from glia to NBs. Similar to full length Hh (Figure 2J-K), glial HhN expression caused a reduction in NB EdU incorporation (Figure 7D), suggesting that palmitoylated Hh is sufficient for glia-NB crosstalk.

Similar to Rasp knockdown, Fasn1 knockdown caused a reduction in the number of HhN-GFP puncta that made contact with NBs (Figure 7G-I’, E). To test if Fasn1 is functionally required to mediate HhN’s ability to cause NB S-phase progression defects, we knocked down Fasn1 while overexpressed HhN. In this setting, Fasn1 knockdown rescued EdU incorporation defects caused by HhN expression (Figure 7F). Together, our data suggests that palmitoylation via Rasp and Fasn1, in addition to regulation by Lsd-2, are required for Hh function in the context of glia-NB crosstalk.

### Glial ROS acts in parallel to lipid-Hh signaling to regulate NB proliferation in the FGF-driven glioma model

Lipid metabolism alteration in glia has previously been linked to reactive oxygen species (ROS), where excessive ROS production causes LD accumulation, which in turn triggers neurodegeneration (Bailey et al., 2015; Liu et al., 2015). Furthermore, ROS upregulation is known to promote glioblastoma progression (reviewed by Schieber and Chandel, 2014). We therefore tested whether lipid metabolism changes in the FGF-driven glioma model is correlated with ROS levels. Using ROS-inducible *gstD* promoter-*GFP* reporter (Sykiotis and Bohmann, 2008), we detected a 5-fold increase in *gstD-GFP* upon FGF-activation in glial cells (Figure S7A-C). To decipher the effect of ROS manipulation on lipid metabolism, we overexpressed a *Catalase* (Anderson et al., 2005) and a *superoxide dismutase 1* (Martin et al., 2009) in cortex glial cells where FGF was activated, to suppress ROS. And we found the number of LDs, an indicator of lipid metabolism, was not altered upon ROS reduction (Figure S6F-G, J-K). These manipulations were however effective in partially rescuing NB EdU incorporation defects caused by glial FGF activation (Figure S6L) without affecting glial overgrowth (Figure S6D-E, H-I). We conclude that the production of ROS upon FGF activation in the glial niche potentially acts in parallel with lipid-Hh signaling to inhibit NB proliferation.

Collectively, our findings indicate that the expression of Hh in the cortex glia is required for the formation of glial chambers. Moreover, we demonstrate that expression of Hh in the cortex glial niche must be restrained to prevent ectopic Hh signalling in the NBs. We showed that the ability of glial Hh to signal to NBs is modulated by lipid metabolism enzymes Fasn1 and Lsd-2. Upon overgrowth of the cortex glial niche induced by FGF activation, both Hh and lipid metabolism regulators are upregulated in the niche, which in turn slow down the NB cell cycle, and affect its ability to generate a full repertoire of neurons (Figure 7J).

## Discussion

In the mammalian CNS, neuron and glia interactions are complex. Astrocytes that are structurally related to the cortex glia in *Drosophila*, enwrap multiple neurons and NSCs, and are known to modulate adult neurogenesis through soluble signals such as morphogens and extracellular matrix proteins (reviewed by Spampinato et al., 2019). Similarly, the *Drosophila* cortex glial niche, which form chambers that encapsulate NB and its progeny, creates microenvironments that are required for NB maintenance and neuronal maturation (Coutinho-Budd et al., 2017; Dumstrei et al., 2003; Pereanu et al., 2005; Speder and Brand, 2018; Yuan et al., 2020). Furthermore, glial overgrowth observed in the context of glioblastoma has recently been shown to affect NBs and neuronal survival (Portela et al., 2019; Read, 2018). In this study, we have identified a new mechanism of glia-NB crosstalk via Hh modulated by lipid regulators, that affect NB cell cycle progression and lineage size.

Hh has been shown to play pro-proliferative roles in astrocyte proliferation in the mammalian brain (Ugbode et al., 2017), consistent with this, we found Hh is required for the growth of cortex glial cells. During development, NB lineages have been shown to produce its own pool of Hh (Chai et al., 2013), which likely act redundantly with the glial Hh. Hh is however required for glial chamber formation, and Hh knockdown caused a decrease in glial niche, reminiscent of phenotypes reported for glial PI3K inhibition (Speder and Brand, 2018). Furthermore, glial Hh knockdown and other glial niche disruptions caused an increase in M phase NBs and a delay in S phase progression (Speder and Brand, 2018). This contrasts with Hh and Smo mutant NB phenotype, previously reported by Chai et al., 2013, where they found the loss of NB Hh signaling caused a significant increase in NB clone size. Therefore, we favour the model that Hh is required for niche elaboration, which in turn affects NB proliferation.

Glial Hh overexpression (directly and via FGF activation) affected both NBs cell cycle progression and polarity protein localisation (data not shown), reminiscent of the phenotype reported for *ptc* mutant NBs (Chai et al., 2013). Hh closely associates with LDs in the cortex glia. LDs are analogous to lipoproteins, which have been shown to transport Hh for long-range signalling (Palm et al., 2013; Panakova et al., 2005). Different from lipoproteins, LDs mainly act as lipid storage organelles, and are less mobile. Our data supports the model that LD might impede the trafficking of Hh and affect their ability to signal to NBs. Only under Hh overexpression, Hh is capable of signalling to NBs.

Different to other secreted molecules that play a role in the glial niche, such as Dlp (Kanai et al., 2018) and Jeb (Cheng et al., 2011), that are necessary but not sufficient to induce changes in the NB, Hh plays a physiologically relevant role in a disease model for glioma caused by FGFR activation. TCGA data shows that FGFR1–4 are expressed to varying degrees and in different combinations in patient samples, and FGFR1 is an important contributor to poor outcome in glioblastoma (reviewed in Jimenez-Pascual and Siebzehnrubl, 2019). In the *Drosophila* brain, FGF activation caused an expansion of cortex glia where Hh-LD associations are normally observed (Avet-Rochex et al., 2012; Avet-Rochex et al., 2014). This led to an increase in Hh, which in turn affected NB proliferation. In contrast, glial InR and EGFR activation that are insufficient to cause cortex glial overgrowth also did not affect NB proliferation. Together, our data suggests that cortex glia is the key glial subtype that is responsible for glia-NB crosstalk, consistent with the role of cortex glia in niche formation.

Lipid synthesis and LDs have recently been reported to be upregulated in glioma, and are emerging as important biomarkers and metabolic targets in this context (Geng and Guo, 2017; Guo et al., 2013). Here, we found genes that promote lipid synthesis and storage were upregulated in glial overgrowth caused by FGF activation. How does lipid metabolism regulate Hh function? We showed two mechanisms. The first mechanism involves the interaction between lipid storage regulator Lsd-2 and Hh that colocalize on the surface of LDs and that Lsd-2 modulates Hh’s ability to signal to NBs. It is not yet clear how Lsd-2 directly regulates Hh activity, but we think it is possible that Lsd-2 might physically interact with Hh and affect its secretion; or alternatively, Lsd-2 competes with Hh for positions on the surface of LDs, and pushes Hh into the cytoplasm. The second mechanism involves Fasn1and Rasp which regulates Hh-post translational modification. In addition to the Hh-LD axis, ROS which is implicated in glioblastoma (Schieber and Chandel, 2014) also lies downstream of FGF mediated glia-NB crosstalk. Knockdown of both Hh and ROS-axes significantly rescued NB cell proliferation defects caused by FGF activation in the glial cells. The phenomenon we reported here together with other reports that glioblastoma affects the survival or proliferation rate of its neighbours (Portela et al., 2019; Read, 2018) poses an interesting but yet unexplored prospect that glioma outcompetes NBs within the CNS for limited energy and nutrient resources.

## Acknowledgements

We are grateful to Alex Gould, Thomas B. Kornberg, Isabel Guerrero, Yu Cai, William Chia, Tatsushi Igaki, Joseph M. Bateman, Kieran Harvey, Helena Richardson, Gary Hime, Philip Batterham for generous sharing of antibodies and fly stocks. We would like to thank Bloomington Drosophila Stock Center, Vienna Drosophila Resource Center, Fly Stocks of National Institute of Genetics, KYOTO Stock Center, Developmental Studies Hybridoma Bank and Addgene for fly stocks and plasmids. We would like to also thank OZDros for *Drosophila* quarantine, Peter MacCallum Cancer Institute Microscopy core and Biological Optical Microscopy platform at the University of Melbourne for technical assistance. We would like to thank Charles Robin, Mike Murray for sharing their microinjection facility with us. We are grateful to Kieran Harvey, Helena Richardson and Andrew Cox for critical reading of the manuscript. Schematic pictures in the figures are created with BioRender. Q.D. is funded by a Melbourne Research Scholarship, L.Y.C is funded by an ARC Future Fellowship, L.Y.C’s laboratory is supported by funding from the NHMRC, ARC and the Peter MacCallum Cancer Foundation.

## Author contribution

Q.D, M.Z, F.F, T.L, S.G conducted the experiments; Q.D and L.Y.C designed the experiments and wrote the paper.

## Method and materials

### Fly husbandry and strains

Fly stocks were reared on standard *Drosophila* media at 25 °C. Crosses for overexpression and knockdown experiment were set up at 25 °C, and after a day, the progenies were moved to 29 °C, unless otherwise stated.

The fly strains used were: *repo-GAL4* (BDSC7415), *NP2222-GAL4* (KYOTO112830), *wrapper-GAL4* (Coutinho-Budd et al., 2017; Richier et al., 2017), *dnab-GAL4* (From Alex Gould lab), *repo-LexA::GAD* (BDSC67096), *w^1118^*, *UAS-htl^ACT^*(BDSC5367), *UAS-Egfr^ACT^*(BDSC59843), *UAS-InR^wt^*(BDSC8262), *LexAop-htl^ACT^* (generated in this paper), *UAS-GFP, UAS-mGFP, UAS-dcr2, UAS-FUCCI* (BDSC55110), *UAS-lacZ*, *UAS-luc* (BDSC64774)*, UAS-hh* (from Thomas B. Kornberg lab), *UAS-hh.N.GFP* (BDSC81023), *UAS-lsd2* (from Alex Gould lab), *UAS-ci^Nc5m5m(ACT)^* (from Yu Cai lab), *UAS-cat.A* (BDSC24621), *UAS-Sod1.A (BDSC24754)*,*UAS-mcherryRNAi* (BDSC35785), *UAS-GFPRNAi* (BSDC9331), *UAS-fasn1RNAis* (NIG3523R-2, NIG3523R-6), *UAS-lsd2RNAis* (VDRC102269, BDSC34617 and BDSC32846), *UAS-hhRNAis* (VDRC43255, BDSC25794), *UAS-ciRNAis* (NIGHMJ23860, NIG2125R-2), *UAS-raspRNAis* (NIG11495R-1, NIG11495R-2), *dpnGFP* (BDSC59755), *his2AV-mRFP* (Kieran Harvey lab), Hh:GFP BAC, Ptc:mcherry BAC*, ci-lacZ* (all three lines are generated by Thomas B. Kornberg lab), *lsd2YFP* (KYOTO115301), *gstD-GFP* (From Tatsushi Igaki lab). The *repo*-MARCM stock was: *UAS-RedStinger; repo-flp, repo-GAL4, UAS-actinGFP; FRT82B, tub-gal80* (from Joseph M. Bateman lab)*. w;;FRT82B* was used to generate surface glial clones.

### Immunostaining

Larval and pupal brains were dissected in PBS, fixed for 25 minutes in 4% formaldehyde in PBS and rinsed in 0.5% PBST (PBS + 0.5% TritonX-100). For immunostaining, brains were incubated with primary antibodies overnight at 4 °C, followed by an overnight secondary antibody incubation at 4 °C. Samples were mounted in 80% glycerol in PBS for image acquisition. The primary antibodies used were mouse anti-Mira (1:50; gift of Alex Gould), rat anti-Mira (1:100, Abcam), rabbit anti-Mira (1:200, gift of Rita Sousa-Nunes), rat anti-pH3 (1:500; Abcam), rat anti-Elav (1:100, DSHB), chick anti-GFP (1:2000; Abcam), rabbit anti-RFP (1:100, Abcam), mouse anti-Fibrillarin (1:200, Abcam), rabbit anti-Hh (1:500, gift of Isabel Guerrero), rabbit anti-Insc (1:20, gift of William Chia). Secondary donkey antibodies conjugated to Alexa 555 and Alexa 647 and goat antibody conjugated to Alexa 488, 555 and 647 (Molecular Probes) were used at 1:500. DAPI (Molecular Probes) was used at 1: 10,000.

### EdU labelling and pulse chase

EdU in vitro labelling was used to identify actively dividing NBs, larval brains were M EdU /PBS for 15 mins (for 96ALH brains) or 1 hr (Figure S4P-S), followed by fixation, and development using Click-iT Plus EdU Cell Proliferation Kit for imaging, Alexa Fluor 647 dye (Invitrogen), following the manufacturer’s instruction. The brain tissues were then stained with Mira to label NBs. In the procedure, control brains and experimental brains were processed in the same tube.

EdU pulse-chase was used to identify the progeny of dividing NBs. 3^rd^ instar larvae were fed with instant fly food supplemented with 100 μg/mL EdU (Lee et al., 2006) for 3 hours. They were then transferred to standard *Drosophila* media for a 4 hour-EdU chase. Wandering stage larvae were collected for brain dissection, followed by fixation, development and immunostaining as described above.

### LD staining

For LD staining, larval brains were dissected in PBS, fixed, and rinsed in PBS before incubation in HCS LipidTOX Red Neutral Lipid Stain (Invitrogen, 1:1000 in PBS) for 1 hour. These samples were then rinsed and mounted in PBS and imaged directly to preserve LD morphology. For experiments that require LD staining together with immuno-staining, the tissues were rinsed three times in PBS after immunostaining to remove all PBST, and incubated with LD dyes as described above. The tissues were mounted in 80% glycerol in PBS for imaging.

### Imaging and image processing

Images were collected on a Leica SP5 or Olympus FV3000 confocal microscopes and analysed using Fiji (https://imagej.net/Fiji). Z stacks of CBs were imaged and a single section of the ventral side of the CB was shown as the representative image unless otherwise stated.

### Live cell imaging

Dissected brains (72ALH) were cultured in Schneider’s culture medium supplemented with 10% inactivated FBS, 2% Penicillin-Streptomycin solution, 20μ glutamine and Schneider’s culture medium and dissected fat body from the same animals. The brains were imaged in a μ-Slide 8 well (Ibidi) on an Olympus FV3000 microscope using resonance scanner in 16Bit mode, with a 40x/0.95 lens and 2x zoom. Z stacks with 2 μm intervals were captured every 2 minutes over a period of 3-8 hours. Laser intensity were kept low to avoid cytotoxicity. AVI movies were generated using Fiji.

All the images were processed using Adobe Photoshop and compiled using Adobe Illustrator.

### Quantitative reverse transcription PCR

For gene expression analysis, 20 dissected late 3^rd^ instar larval brains were lysed in 300 μl TRI Reagent (ZYMO Research R2061) to form one biological replica. Three biological replicates were prepared for each genotype: *repo-GAL4> w^1118^* and *repo-GAL4> htl^ACT^*. Total RNA was extracted using a Direct-zol RNA Microprep Kit (ZYMO Research R2061) and 1μg of total RNA from each sample was reverse transcribed into cDNA using ProtoScript II First Strand cDNA Synthesis Kit (NEB, E6560S) according to the manufacturer’s instructions. The qPCR was performed using the stepOnePlus real-time PCR system (Applied Biosystems) using Fast SYBR Green master mix reagent (Applied Biosystems, 4385612). Gene expression levels were normalized to *rpl32*, and calculated using the 2^-Δ^ method. The primers were either designed using Primer-BLAST (https://www.ncbi.nlm.nih.gov/tools/primer-blast/), or obtained from FlyPrimerBank (https://www.flyrnai.org/flyprimerbank). listed in Supplemental Table 1.

### Molecular cloning

A constitutively active form of *htl*, comprising the dimerization domain of the bacteriophage λ repressor (described in Michelson et al., 1998), was amplified from the genomic DNA of the fly stock: *UAS-htl^ACT^*(BDSC5367), using a forward primer, 5’-CAACTGCAACTACTGAAATCTGCC-3’, and a reverse primer 5’-CCCCC<u>TCTAGA</u>TTAATAATTACACCACTTCTGC-3’. The resulting PCR product was digested with *Not*I (Promega, Cat#R6431) and *Xba*I (Promega, Cat#R6181), which cut at the restriction sites as indicated in the reverse primer (underlined above). The plasmid, *pJFRC19-13XLexAop2-IVS-myr::GFP* (Addgene_26224), was identically digested to remove *myr::GFP*. The restriction fragment, *Not*I-htl^ACT^-*Xba*I, was subsequently cloned into the digested LexAop vector (Pfeiffer et al., 2010). The reconstructed plasmid was sequenced and injected into flies carrying an attP2 docking site (BL25710). The overexpression of *LexAop-htl^ACT^* using *repo-LexA* recapitulated the glial overgrowth caused by *UAS-htl^ACT^* overexpression using *repo-GAL4* (data not shown).

## Quantification and analysis

### Cortex glial membrane size and number measurement

Cortex glial membrane volume (*NP2222-GAL4>UAS-GFP*) were measured from three-dimensional reconstruction of confocal Z stacks (2-μm step-size) with Volocity software (Improvision). Glial cell number were automatically counted with a Fiji plug-in “DeadEasy Larval Glia” (Forero et al., 2012).

### Cell cycle speed

**The number of Type I NBs** in each brain lobe was manually counted using Fiji. Type I NBs were distinguished from other Mira^+^ cells by size and morphology: Newly generated GMCs are smaller than Type I NBs as shown in Figure S3D (white arrows). Type II NBs are associated with more Mira^+^ progeny cells.

**EdU index:** For 26ALH larval brains, EdU voxels of the whole brain were measured with Volocity software (Improvision) to indicate NB re-entry into cell cycle. Glial EdU voxels represents only a small amount of the total EdU voxels (Sousa-Nunes et al., 2011). Normalized EdU voxels were calculated by dividing EdU voxels to the mean voxel counts of the control. For 96ALH larval brains, the number of EdU^+^ Type I NBs was counted in the central brain of each brain lobe. EdU index was determined by normalizing the number of EdU^+^ Type I NBs to the average number of EdU^+^ Type I NBs in the control. The total number of Type I NBs is not altered in experiments where Edu index was used to determine NB cell cycle speed. For EdU pulse-chase quantification, the number of neurons that inherit EdU from each dividing NBs was counted to indicate the speed by which NBs generate their progeny.

**pH3 index:** represented as the % of Type I NBs in M phase (pH3^+^)/ the total number of Type I NBs.

**Cell cycle lengths in NBs** were measured as the time between two consecutive cell divisions. The cell cycle lengths of NBs from at least three different ex vivo brains were plotted for each genotype.

### Cell size measurement

NB, GMC and nucleoli diameters were estimated by averaging orthogonal measurements of diameter, with a single confocal section on Fiji.

**Localization of asymmetric determinants** was assessed in M phase NBs that display a condensed metaphase plate marked by pH3. Clear crescent localisation was counted as correct localisation, and cytoplasmic or cortical localisation was counted as mislocalisation.

**The distribution of Hh in glia** was measured on Fiji with the formula: The area of glia that contains Hh / Total glial area. The detailed procedure: Total glial area: Adjust Threshold (Default) > Analysis > measure (area); The area of glia that contains Hh: Create selection of glial channel by Adjusting Threshold (Default) > Restore selection in the Hh channel > clear outside > measure area.

### Total GFP intensity

The single confocal image was used for the intensity measurement on Fiji using the formula: CTCF (corrected total cell fluorescence) = Integrated density – (Area of selected cell x mean fluorescence of background readings) (McCloy et al., 2014).

### Statistical analysis

*P-*values are calculated by two-tailed, unpaired Student’s *t*-test, with equal sample variance; The Welch’s correction was applied in case of unequal variances. Mann-Whitney test was used when data deviated from a normal distribution. For all histograms, error bars represent SEM.

---

**Supplemental Table 1.**
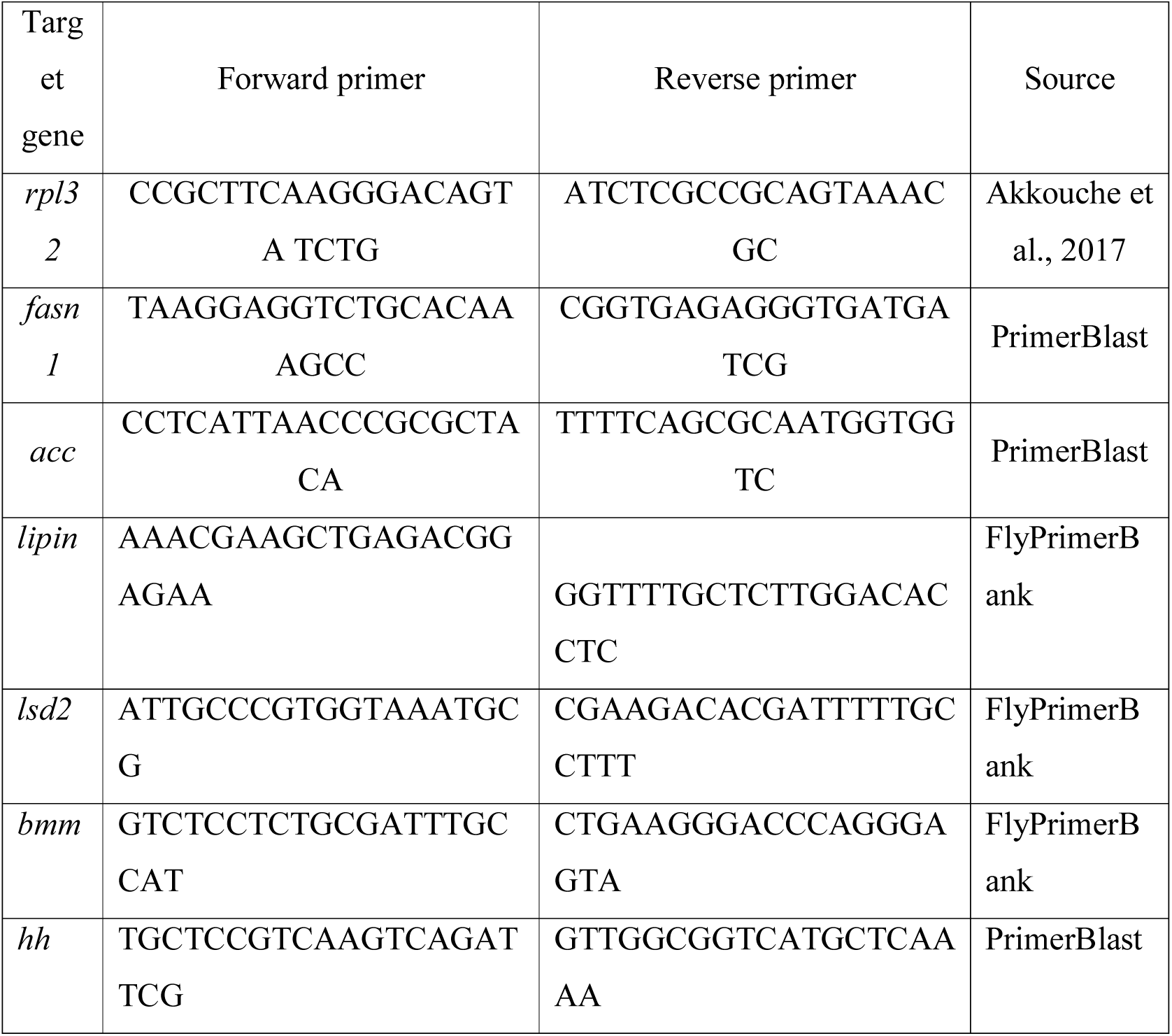

**Figure S1:**
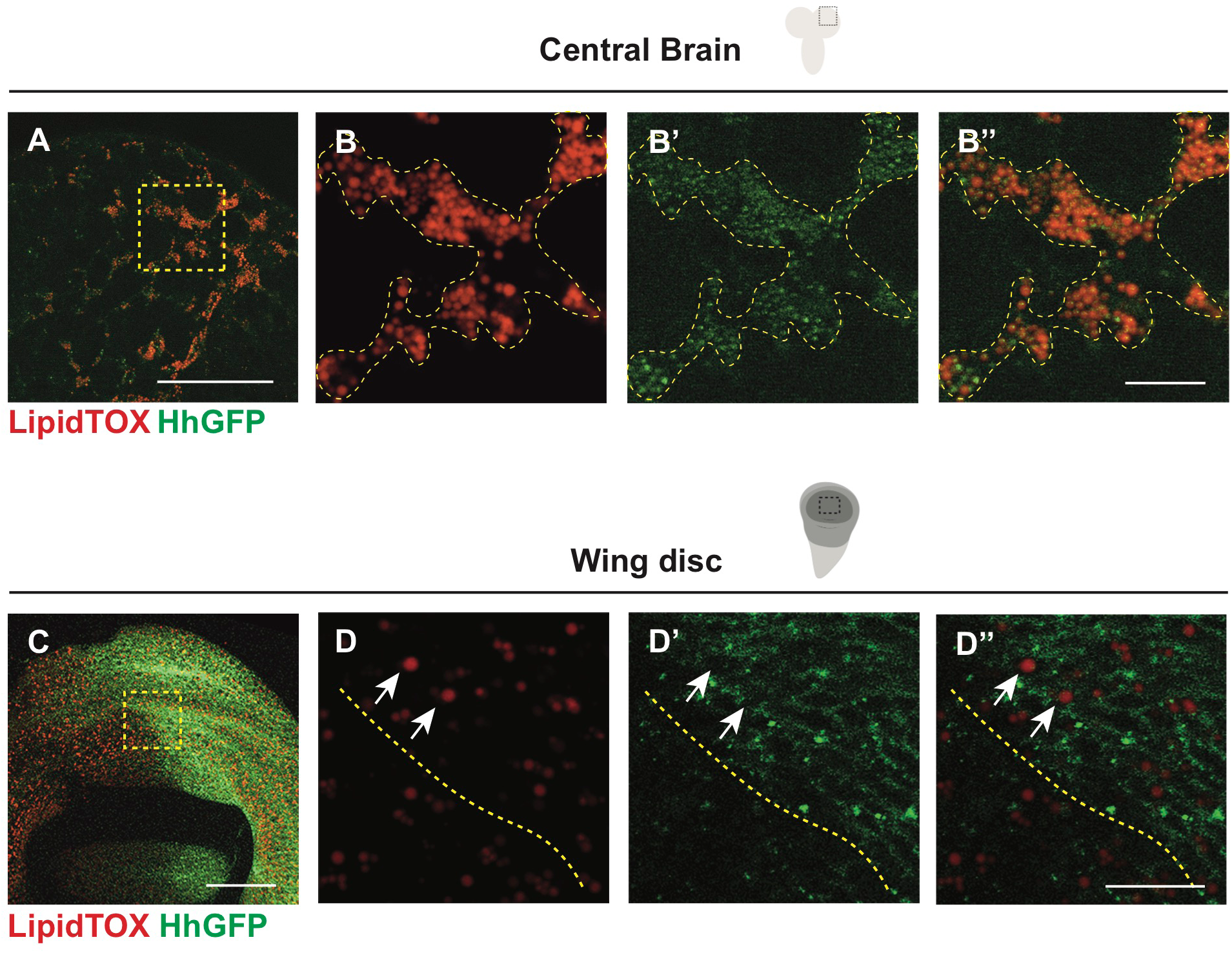
Hh forms complexes with LDs in the CB but not the wing discs (related Figure 1) A-B’’) HhGFP and LDs are associated in the CB glial cells (outlined with yellow dashed lines). C-D’’) HhGFP is not associated with LDs in the posterior wing disc (white arrows). B-B’’, D-D’’ are zoomed in images of A and C, Scale bar= respectively.10 µm in B-B’’ and D-D’’.

**Figure S2:**
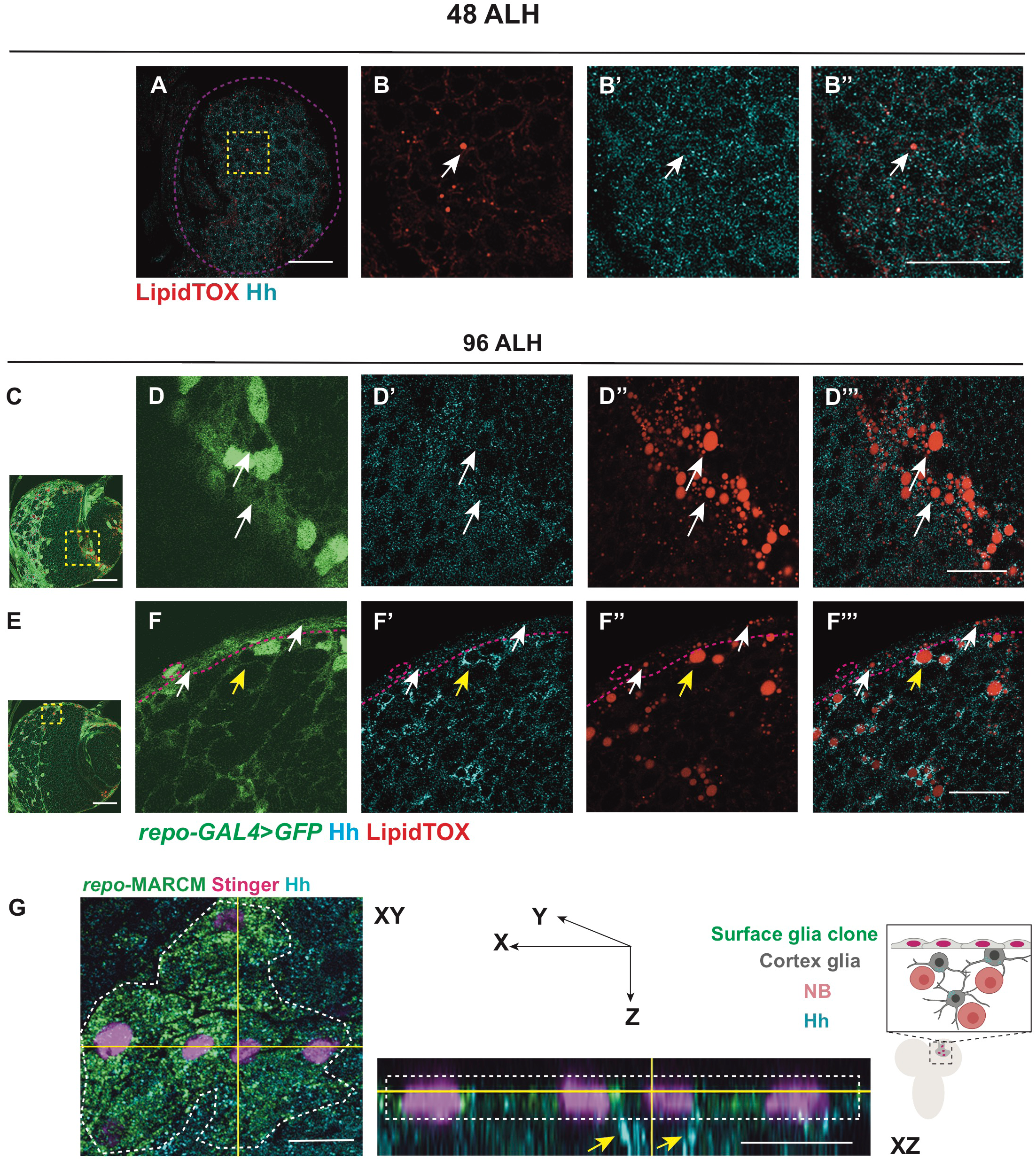
Hh-LD associations are specifically observed in cortex glial cells in the CB during late larval stages (related to Figure 1) A-B’’) Hh and LDs are present at low levels at 48 ALH, and do not form specific association (white arrows). C-D’’’) Hh and LDs are not associated in the optic lobe glial cells (white arrows). E-F’’’) Hh and LDs associate only in the cortex glial cells (yellow arrows) but not the surface glial cells (white arrows). Glial cells are visualised with *repo-GAL4>GFP* in C-F’’’. G) Left and middle panel, representative image showing a surface glial clone (*repo*-MARCM, glial nucleus marked by Stinger in pink). Hh is localised to the cortex glial cells (yellow arrows) underneath the clone marked in green. Right panel, a schematic depicting XZ cross-section of CB glial cells and their relative position. B-B’’, D-D’’’, F-F’’’ are zoomed in images of A, C, E, respectively. Scale bar=20 µm in A-B’’, D-D’’’, F-F’’’, Scale bar=10 µm for XZ section in G.

**Figure S3:**
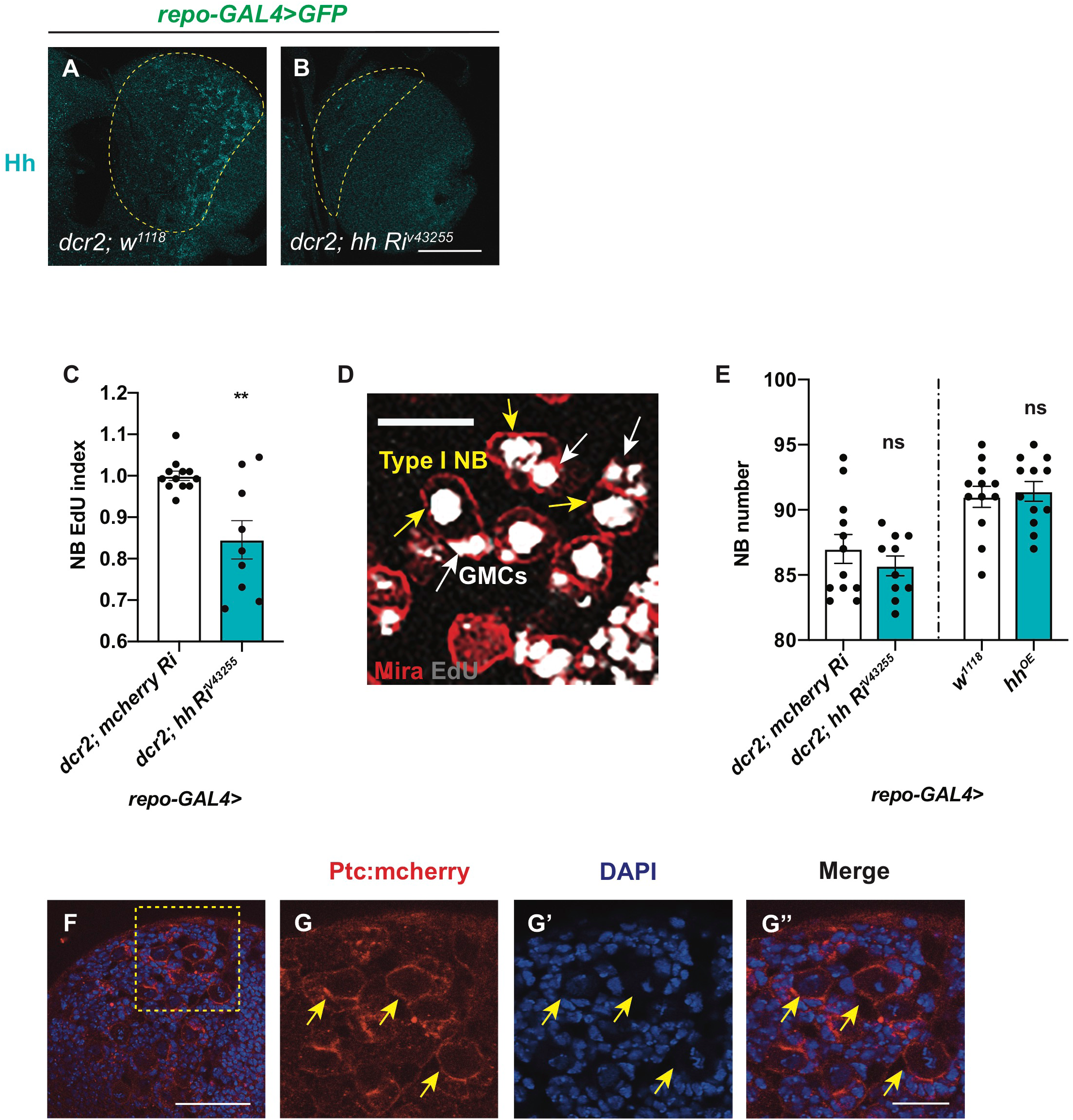
Efects of Hh overexpression and knockdown on Hh level, NB number and EdU index (related to Figure 2) A-B) Representative images showing pan-glial Hh knockdown (*repo-GAL4>GFP* with *UAS-dcr2*) efficiently reduces Hh staining in the CB (outlined with yellow dashed lines). Hh knockdown (*repo-GAL4>GFP* with *UAS-dcr2*) significantly reduces NB EdU index. Representative image from EdU incorporation assays used throughout the manuscript. During a 15 min EdU pulse, type I NB (yellow arrow) and its GMC (white arrow) both incorporate EdU. EdU index quantifications include only EdU^+^ type I NBs. Hh knockdown or overexpression in glial cells (*repo-Gal4>*) does not significantly alter the number of CB NBs. F-G’’) Representative images showing Ptc:mcherry is expressed in NBs (yellow arrows). G-G’’ are zoomed in images of F. Scale bar=20 µm in D, G-G’’.

**Figure S4.**
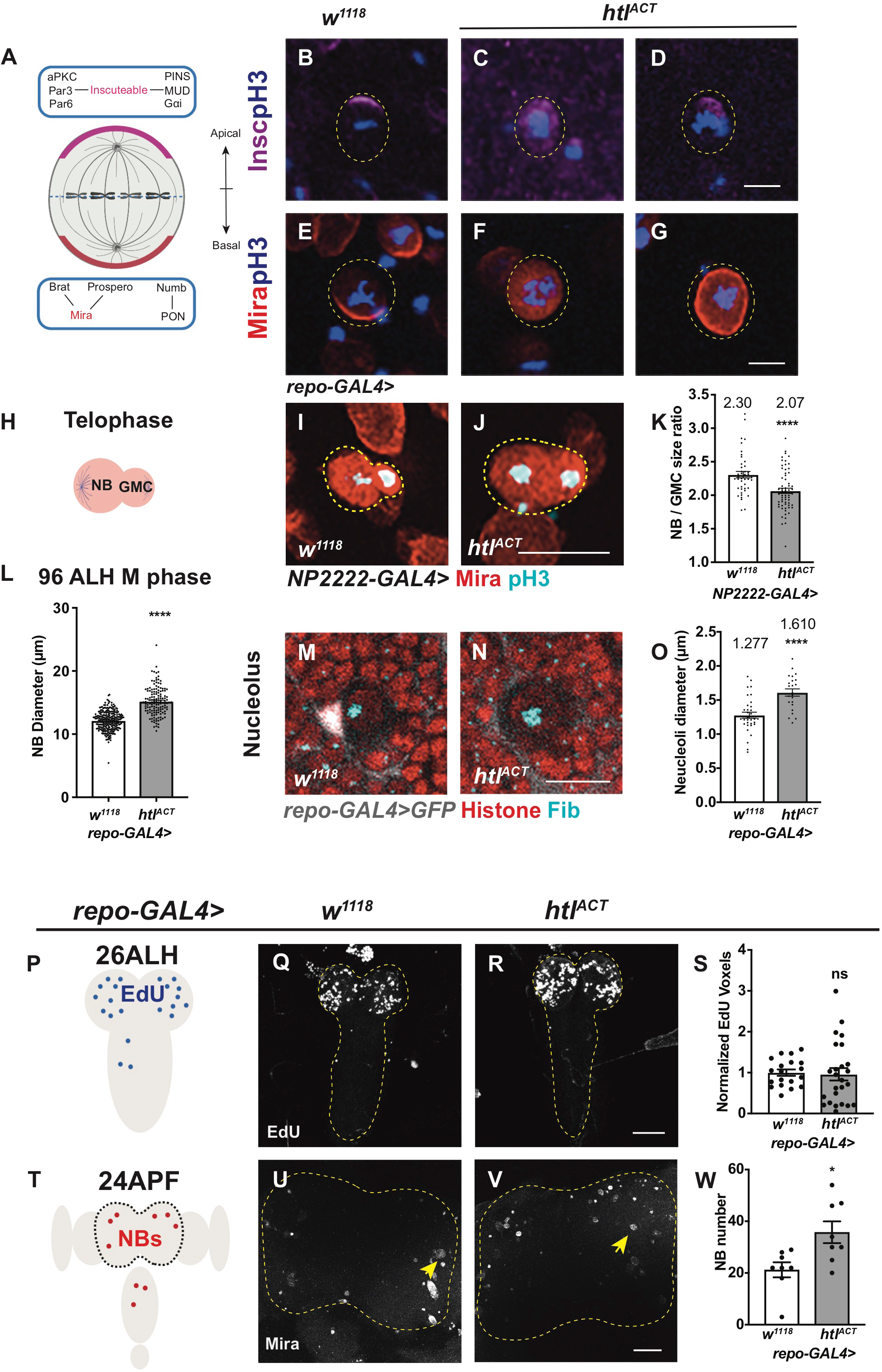
Glial *htl^ACT^* overexpression affects NB asymmetric division, size and cell cycle exit (related to Figure 5) A) Schematic depicting the distribution of polarity proteins in M phase NBs. Apical polarity proteins include the Par complex (aPKC/Par3/Par6), the PINS/MUD/Gαi complex and the adaptor protein, Inscuteable (magenta); Basal polarity complex comprises the cell fate determinants Brat/ Pros/ Numb and their adaptor proteins Mira (red) and PON. B-G) Representative images showing that in pH3^+^ NBs, Insc and Mira mislocalize to the cytoplasm or cortex upon FGF activation in glia (*repo-GAL4> htl^ACT^*). H) Schematic depicting a NB undergoing telophase. I-K) Representative images showing that NBs in telophase (Mira^+^; pH3^+^) give rise to more size-symmetric daughter cells upon cortex glial (*NP2222-GAL4>*) *htl^ACT^* overexpression, quantified in K. NBs are outlined with yellow dashed lines in B-G, I-J. L) Glial (*repo-GAL4>*) *htl^ACT^* overexpression causes an increase in M phase NB diameter. M-O) Representative images showing that NB nucleoli are significantly enlarged upon glial (*repo-GAL4>*) *htl^ACT^* overexpression, quantified in O. NBs are marked by Histone (red), surrounded by glial cells (grey, *repo-GAL4>GFP*), nucleoli are marked by Fib (Cyan). P-S) Representative images showing that the timing of NB cell cycle entry (visualised by EdU incorporation at 26ALH) is not significantly altered by pan-glial (*repo-GAL4>*) *htl^ACT^* overexpression, quantified in S, where EdU voxels are normalized to control. The region of interest is outlined in yellow. T-W) Representative images showing that the number of CB NBs (Mira^+^) at 24APF is significantly increased with pan-glial (*repo-GAL4>*) *htl^ACT^* overexpression, quantified in W. The region of interest is outlined by yellow dashed lines and NBs are marked with yellow arrows. Scale bar=10 µm in B-G; Scale bar=20 mm in I-J and M-N.

**Figure S5.**
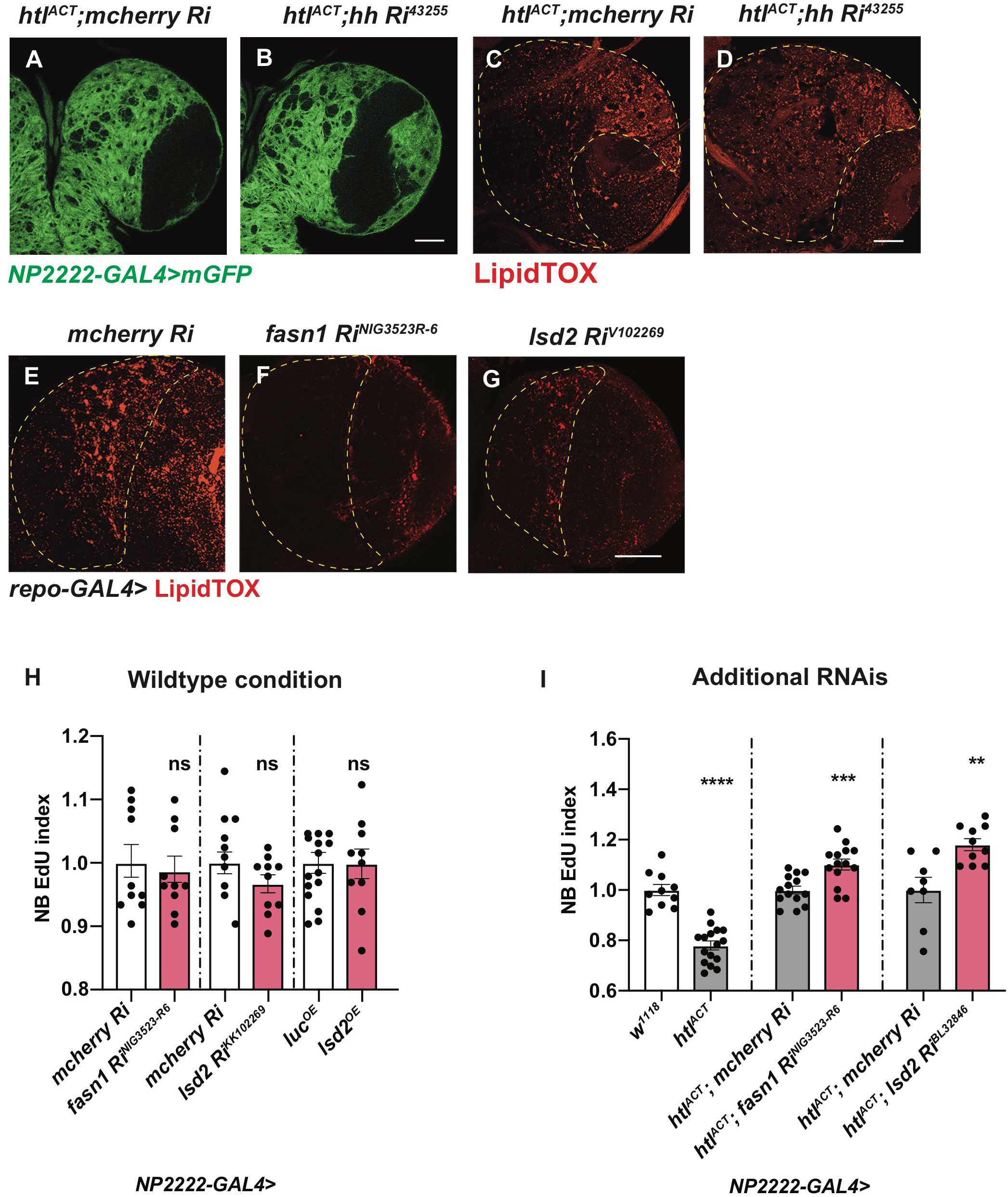
Characterisation of the effects of glial *hh, fasn1* and *lsd2* RNAis on glial size, LDs and NB proliferation (related to Figure 6) A-D) Representative images showing that induction of *hh RNAi* in cortex glial cells with *htl^ACT^* overexpression does not alter the size of cortex glial membrane (*NP2222-GAL4>mGFP*) nor the number of LDs in CB (outlined by yellow dashed lines). E-G) Representative images showing that glial (*repo-GAL4>*) induction of RNAis against *fasn1* and *lsd2* efficiently reduce the number of LDs in CB (outlined by yellow dashed lines). H) knockdown of lipogenesis genes *fasn1* and *lsd2* or overexpression of *lsd2* using a cortex glial driver (*NP2222-GAL4>*) does not significantly affect NB EdU index. I) The NB EdU incorporation defects due to cortex glial (*NP2222-GAL4*) overexpression of *htl^ACT^* is rescued by overexpression of additional RNAi lines against *fasn1* and *lsd2* (related to Figure 6J). The *NP2222-GAL4> w^1118^* vs *htl^ACT^* columns depict the same data as those in Figure 5 E.

**Figure S6.**
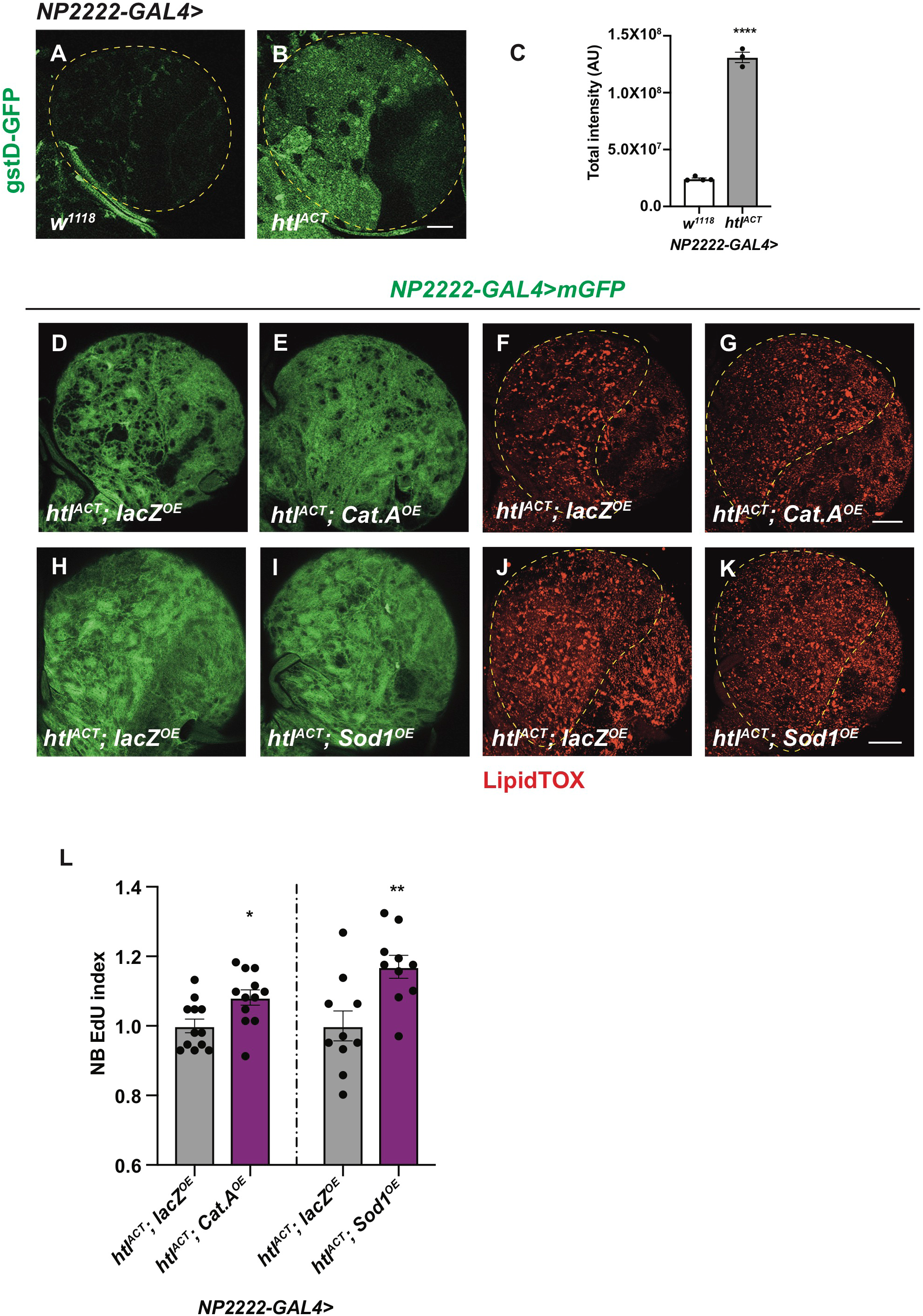
Glial ROS acts in parallel with lipid-Hh signalling to regulate NB proliferation downstream of glial FGF activation (related to Figure 7) A-C) Representative images showing that the oxidative stress indicator gstD-GFP is significantly increased, by cortex glial (*NP2222-GAL4*) overexpression of *htl^ACT^*, quantified in C. Region of interest is outlined with yellow dashed lines. D-L) Representative images showing that cortex glial (*NP2222-GAL4>*) overexpression of ROS scavengers *Cat.A* or *Sod.1* does not alter the size of cortex glia (marked with mGFP) or the number of LDs, but significantly rescues NB EdU incorporation defects caused by *htl^ACT^* overexpression, quantified in L. Region of interest are outlined with yellow dashed line.

